## Supplementary Data for "A cyclic peptide toolkit reveals mechanistic principles of peptidylarginine deiminase IV (PADI4) regulation"

A

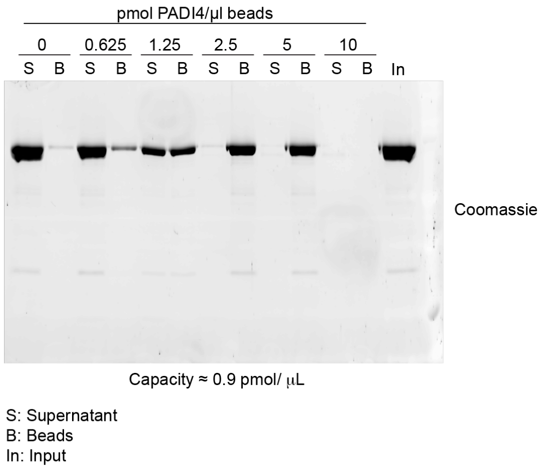

B

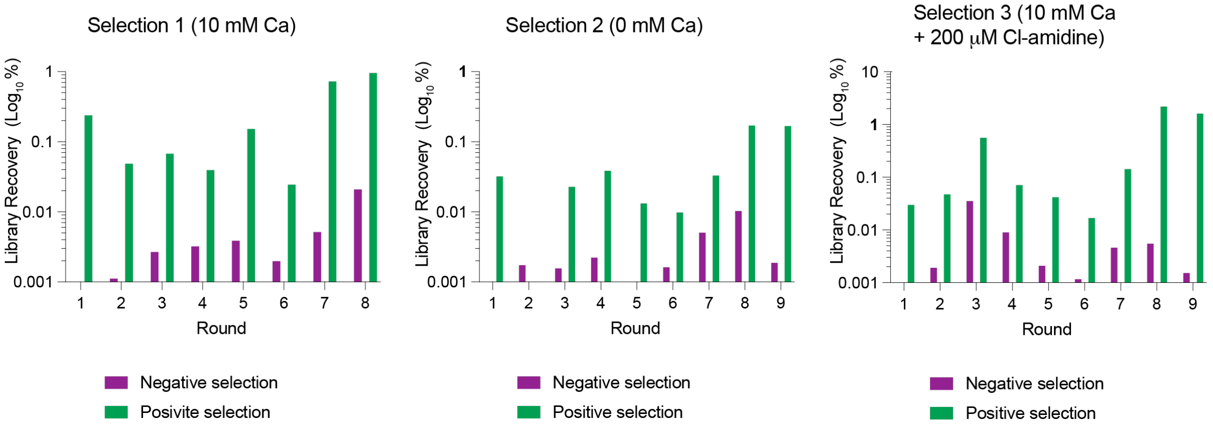

**Supplementary Figure 1**. **RaPID selections with hPADI4. A.** Determination of Avi-His-PADI4 loading level on streptavidin beads. **B.** RaPID selection recoveries. Violet bars and green bars represent the percentage of the input RaPID library recovered after affinity panning against streptavidin beads or streptavidin beads coated with biotinylated PADI4 respectively.

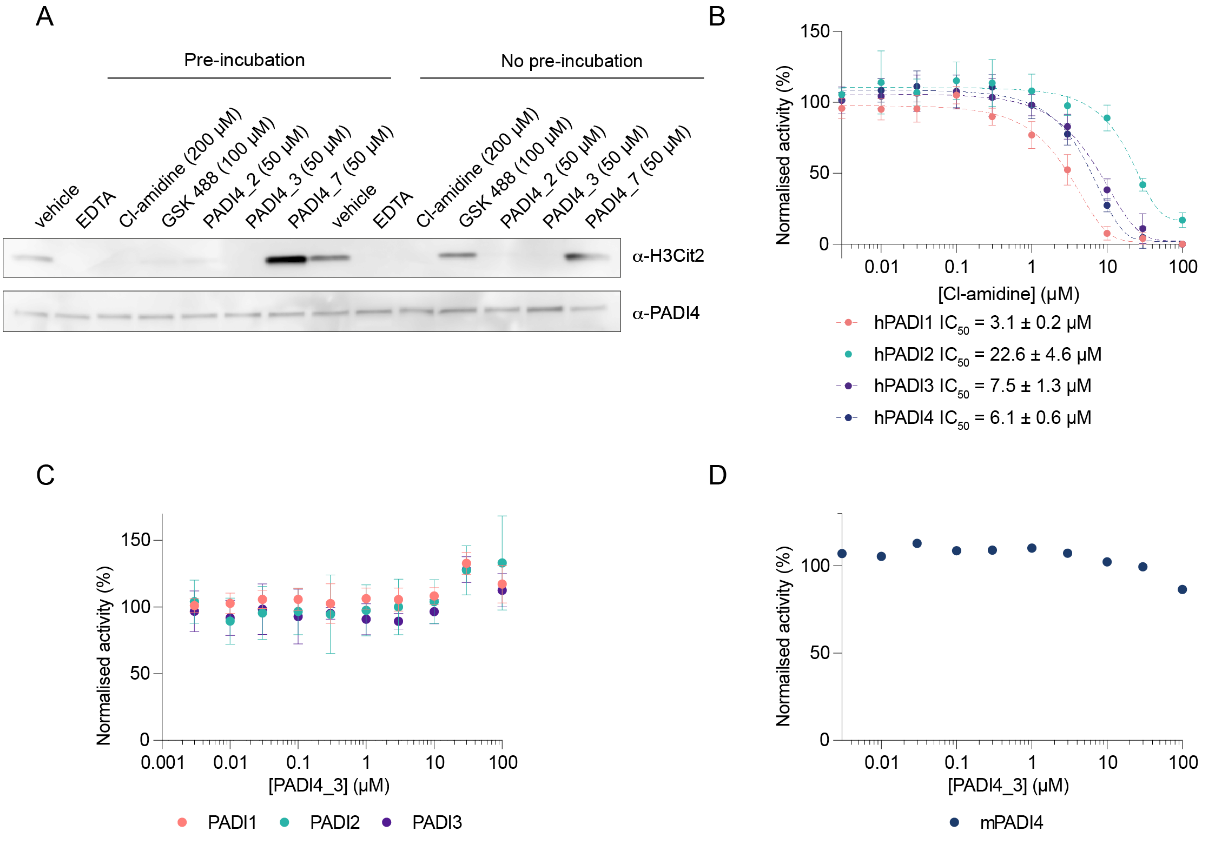

**Supplementary Figure 2. PADI4_3 is a potent and selective hPADI4 inhibitor**. **A.** Immunoblot analysis for citrullinated R2 on histone H3 (H3Cit2), as a measure of PADI4 activity, in hPADI4-stable mES cells. Cell extracts were incubated with indicated PADI inhibitors either 20 mins prior (pre-incubation) or concurrent with the addition of calcium (no pre-incubation). Cellular PADI4 serves as a loading control. **B.** Inhibition of hPADIs by Cl-amidine. COLDER assays of PADI family members with different concentrations of Cl-amidine and 10 mM $CaCl_2$. Data is normalised to activity of each PADI in the presence of 0.1% DMSO vehicle. Data shows mean ± SEM of three independent replicates. Each independent replicate was performed in triplicate. **C.** PADI4_3 does not inhibit PADI1, PADI2 or PADI3. COLDER assays with PADI family members with different concentrations of PADI4_3 and 10 mM $CaCl_2$. Data is normalised to activity of each PADI in the presence of 0.1% DMSO vehicle. Data shows mean ± SEM of three independent replicates. Each independent replicate was performed in triplicate. **D.** COLDER assay of mPADI4 with different concentrations of PADI4_3 and 10 mM $CaCl_2$. Data shows mean ± SEM of three independent replicates. Each independent replicate was performed in triplicate.

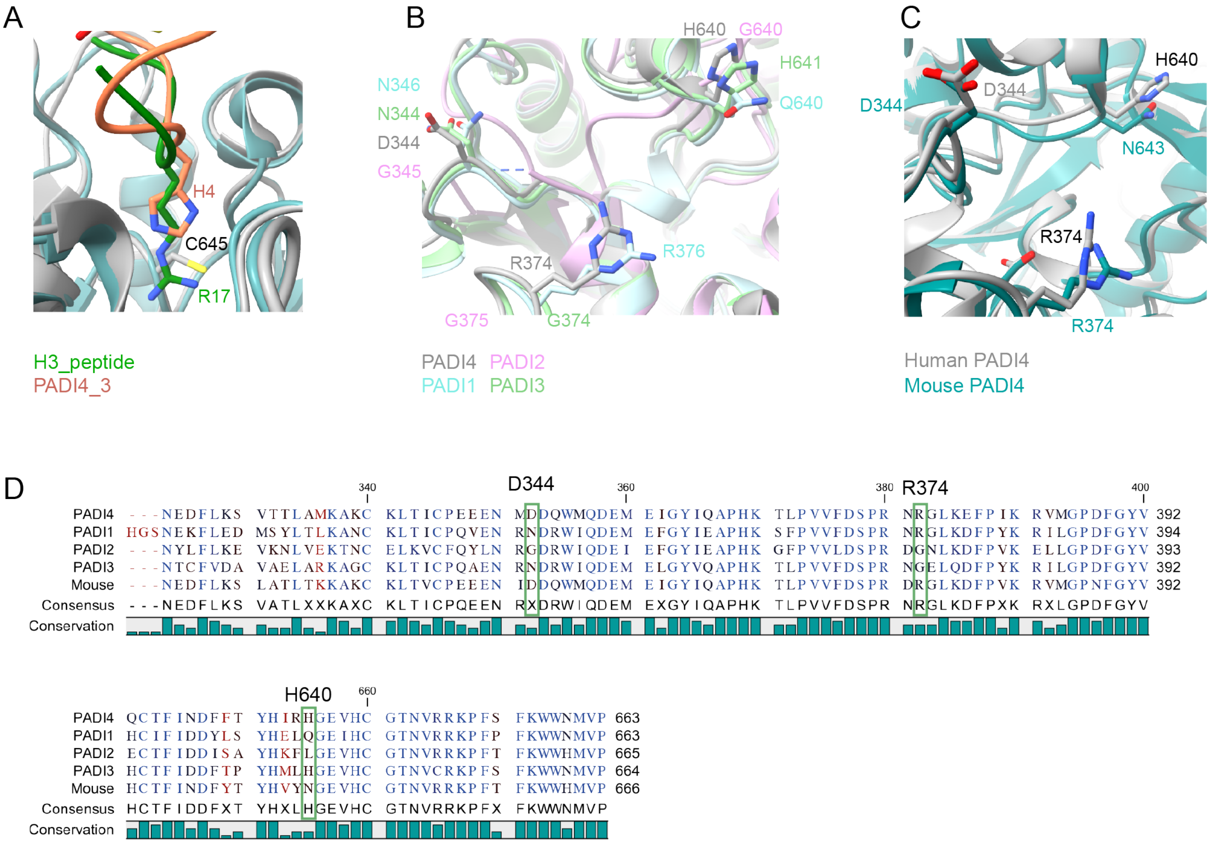

**Supplementary Figure 3**. **PADI4_3 binds to the active site of PADI4 and is selective for human PADI4. A.** Alignment of cryoEM structure of PADI4 bound to PADI4_3 with X-ray crystal structure of PADI4 bound to an H3 peptide with R17 occupying the active site of PADI4 (PDB: 2DEX). PADI4_3 H4 binds in the same pocket as R17. **B.** Alignment of human PADI4 (grey) with other human PADI isozymes. PADI1: AlphaFold model (cyan), PADI2 PDB: 4N2B (pink), PADI3: AlphaFold model (green). Side chains of residues important for interactions with PADI4_3 are shown as sticks. **C.** Alignment of human PADI4 (grey) with its mouse ortholog (teal) (mouse PADI4: AlphaFold model). Side chains of residues important for interactions with PADI4_3 are shown as sticks. **D.** Sequence alignment of human PADI4 with other human PADI enzymes and mouse PADI4. Residues highlighted in green boxes are involved in the binding of PADI4_3 peptide and are not conserved.

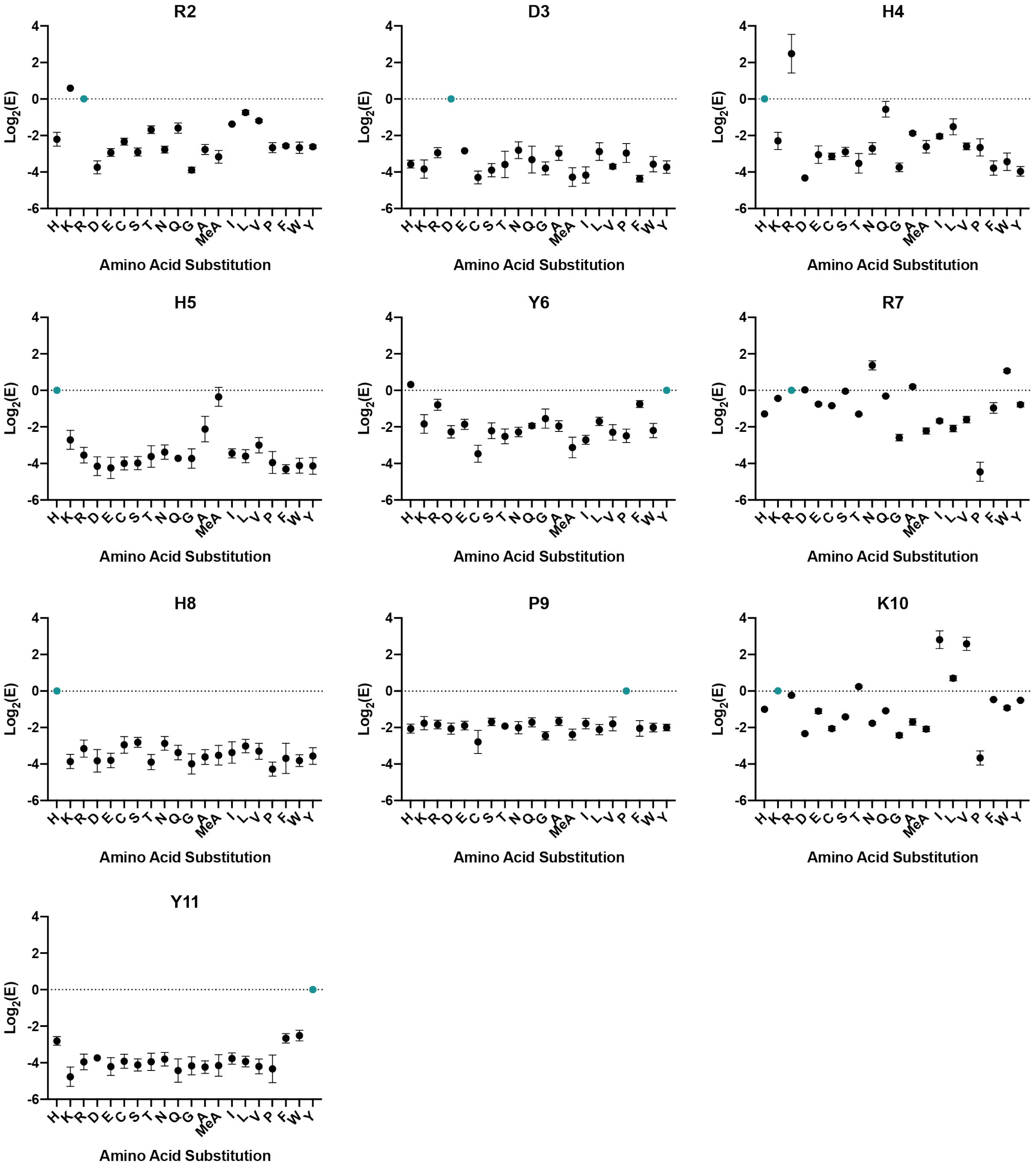

**Supplementary Figure 4. Deep mutational scanning of PADI4_3.** $Log_2(E)$ scores are plotted for each amino acid substitution, for every position varied in the PADI4_3 sequence. Data are plotted as the mean ± standard deviation of three independent single round selections against immobilised PADI4.

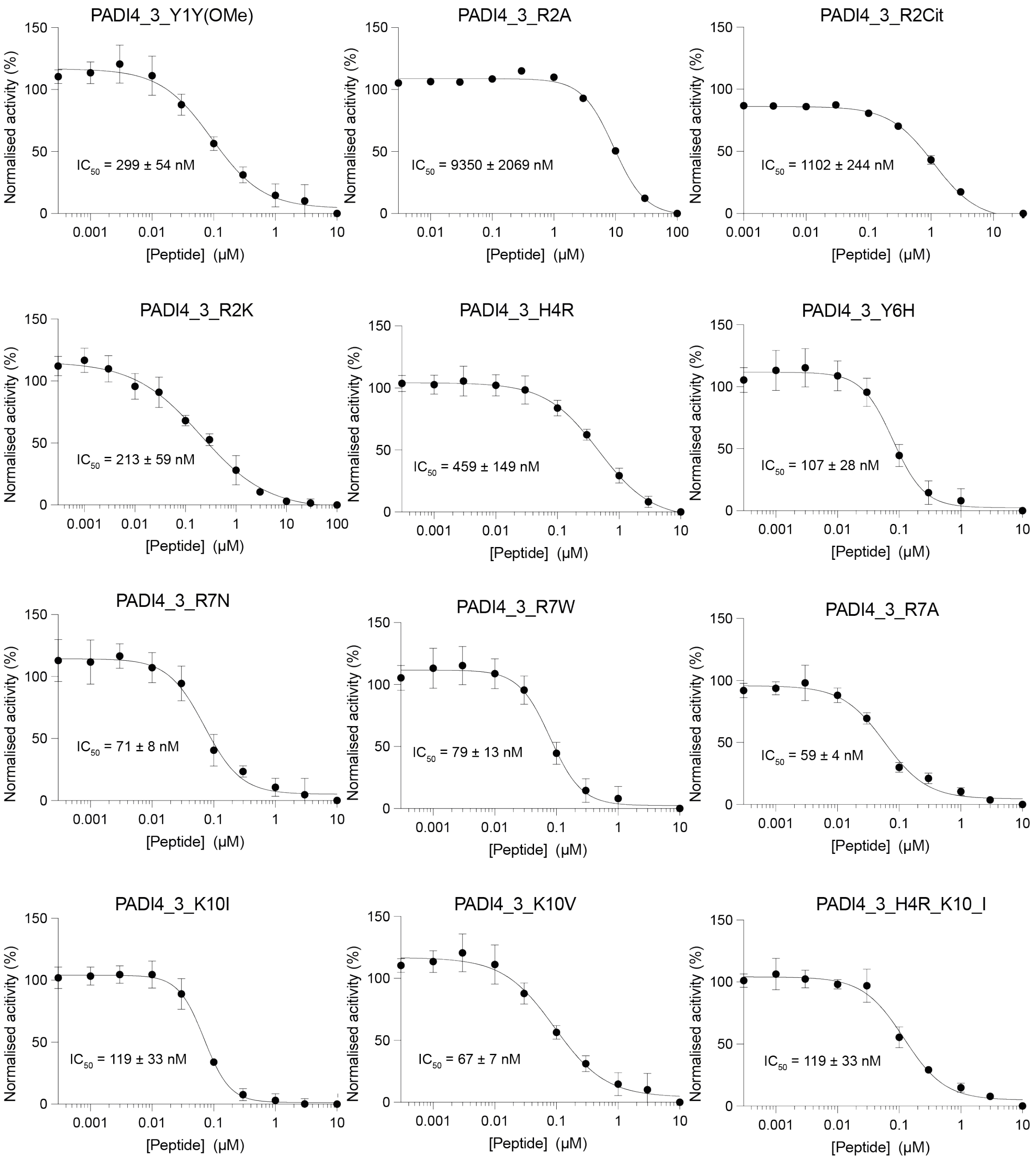

**Supplementary Figure 5**. **Inhibition of hPADI4 by PADI4_3 analogues.** hPADI4 inhibition by different concentrations of peptides (10-0.0003 μM) and 10 mM $CaCl_2$ measured by COLDER assay. Data is normalised to activity of PADI4 in the presence of 0.1% DMSO vehicle. Data shows mean ± SEM of three independent replicates. Each independent replicate was performed in triplicate.

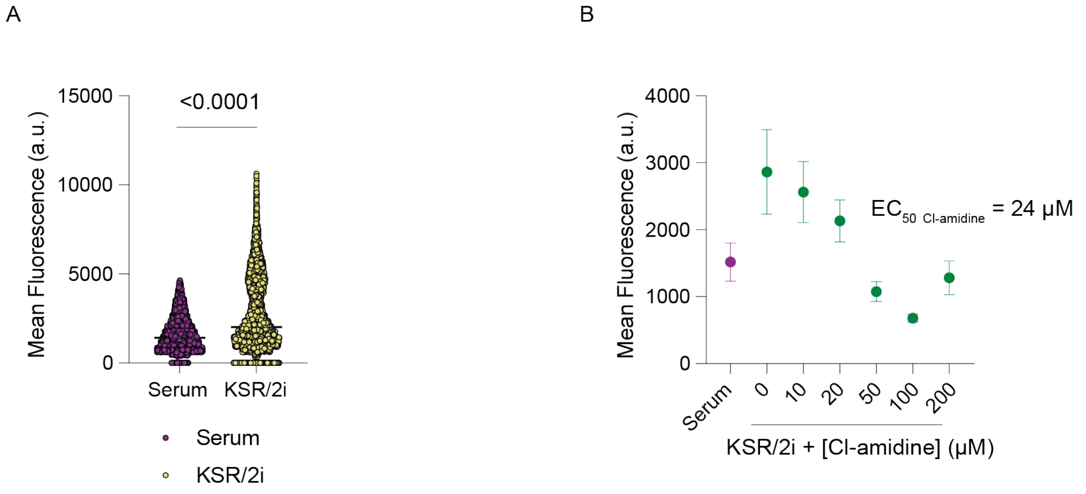

**Supplementary Figure 6. Validation of high content microscopy method.**

**A.** High content imaging-based quantification of mean H3Cit immunofluorescence intensity in hPADI4-stable mES cells grown in Serum or KSR/2i for 3 h. Each data point represents the mean H3Cit intensity per cell. Cells from three technical replicates and three biological replicates are included. **B.** High content imaging-based quantification of mean H3Cit immunofluorescence intensity in hPADI4-stable mES cells stimulated with KSR/2i for 3 h, in the presence of increasing concentrations of Cl-amidine. Each data point represents the mean H3Cit intensity per conditions from three technical replicates.

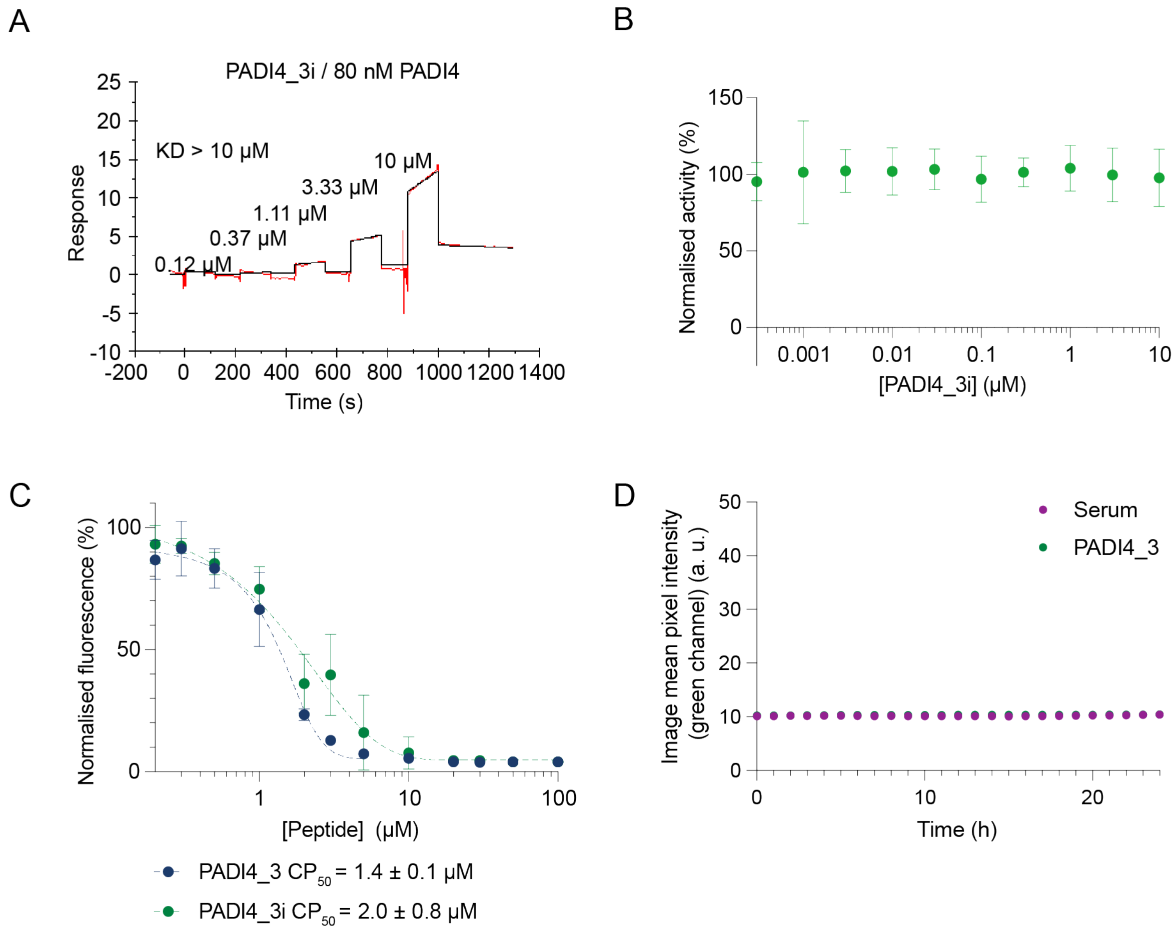

**Supplementary Figure 7. PADI4_3i does not inhibit PADI4 and PADI4_3 and PADI4_3i are cell permeable and non-toxic. A.** PADI4_3i does not bind to PADI4. Binding kinetics between PADI4 and PADI4_3i measured by SPR. A representative experiment is shown. This experiment was performed three times with similar results. **B.** PADI4_3i does not inhibit PADI4. COLDER assay with PADI4_3i at different concentrations (10 - 0.0003 μM) and 10 mM $CaCl_2$. Data are normalised to activity of PADI4 in the presence of 0.1% DMSO vehicle. Data show mean ± SEM of three independent replicates. Each replicate was performed in triplicate. **C.** PADI4_3 and PADI4_3i enter cells. CAPA assay with PADI4_3 and PADI4_3i. Data show mean ± SEM of three different experiments. Each replicate was performed in triplicate. Data are normalised to cells with no peptide treated with TMR dye (Promega) (100 %) and cells with no peptide and no dye (0 %). **D.** Assessment of cytotoxicity by live cell imaging with Incucyte® Cytotox Green Dye, as a measure for cell death. hPADI4 expressing mES cells treated with 1 μM PADI4_3 or DMSO vehicle (0.1%), and imaged for 24h.

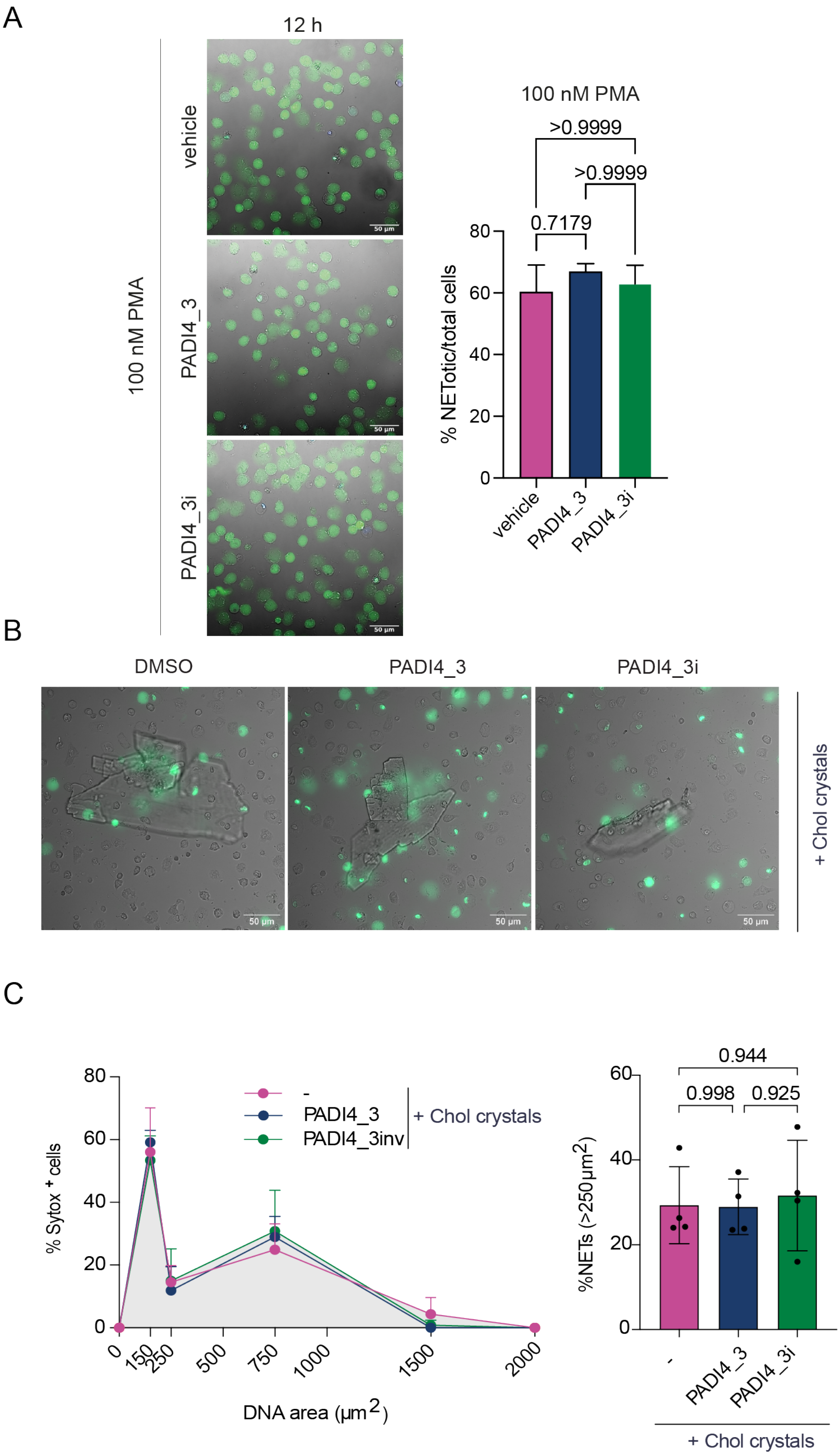

**Supplementary Figure 8. PADI4_3 does not affect NETosis in human neutrophils stimulated with PMA or cholesterol crystals. A.** Bright field micrograph of human neutrophils pre-incubated with vehicle, PAD4_3 or PAD4_3i peptides (50 μM), imaged 12h post-stimulation with PMA (left) and the corresponding quantification of the percentage of

NETotic cells over total cells (right). **B.** Human neutrophils were pre-incubated with DMSO vehicle (0.5%) or 50 µM PAD4_3 or PAD4_3i and the membrane-impermeable dye Sytox-green before stimulation with 0.1 mg/ml cholesterol crystals. Images were acquired by time-lapse fluoresence microscopy, with images at 8h shown. **C.** Quantification of (B). (left) DNA area was measured by Sytox-green signal and areas were distributed into bins of increasing area sizes and plotted as a percentage of Sytox-green positive (dead) cells. Data shown are means +/- SD from one representative experiment. (right) Percentage of NETotic cells in each treatment condition.

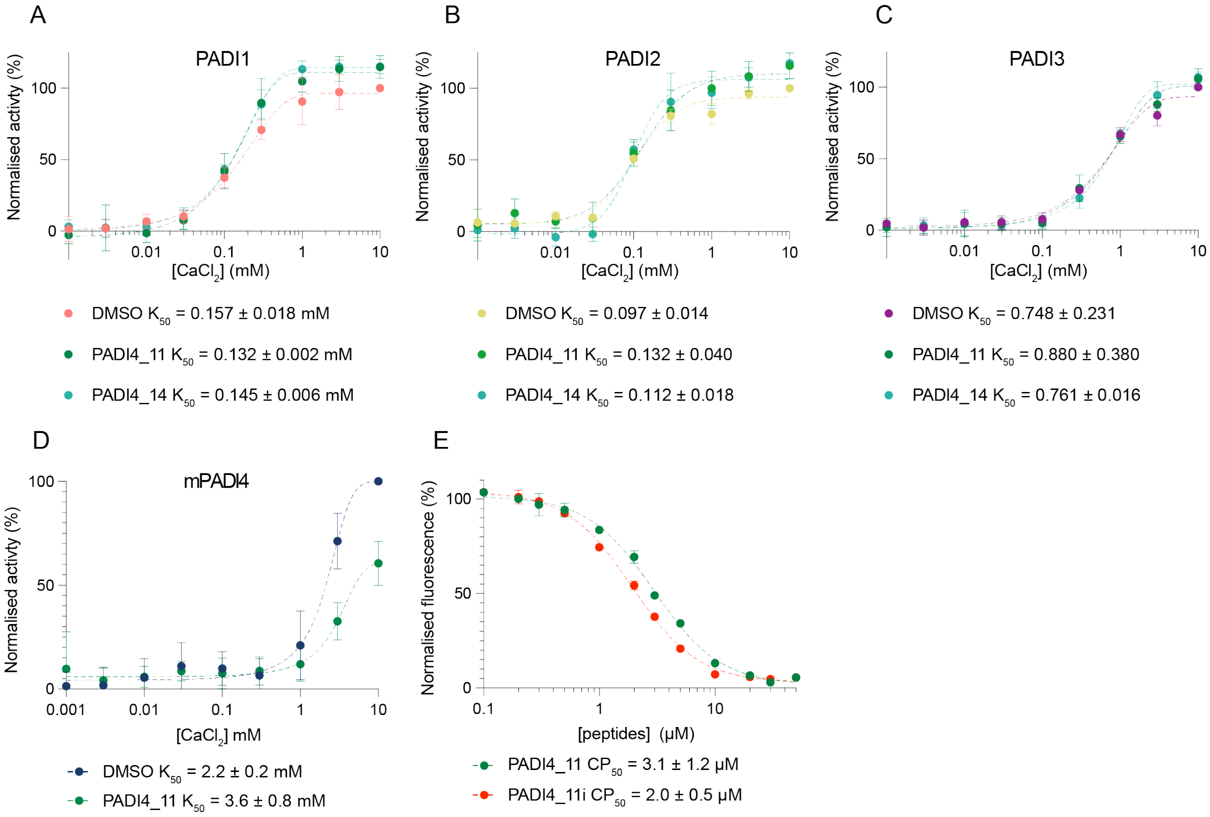

**Supplementary Figure 9**. **PADI4_11 is selective for hPADI4 and is cell permeable. A-D.** PADI4_11 and PADI4_14 do not activate hPADI1 (A), hPADI2 (B), hPADI3 (C) or mPADI4 (D). COLDER assays were performed in presence or absence of PADI4_11 or PADI4_14 at 30 μM and different concentrations of $CaCl_2$. $K_{50Ca2+}$ is the concentration of $CaCl_2$ that yields half maximal PADI activity. Data represent mean ± SEM of three independent replicates. Each replicate was performed in triplicate. Data were normalised against the activity of each PADI in the presence of 0.1% DMSO vehicle and 10 mM $CaCl_2$. **E.** PADI4_11 and PADI4_11i are cell permeable. CAPA assay with PADI4_11 and PADI4_11i. Data show mean ± SEM of three independent experiments performed in triplicate. Data are normalised with cells with no peptide treated with TMR dye (Promega) (100 %) and cells with no peptide and no dye (0 %).

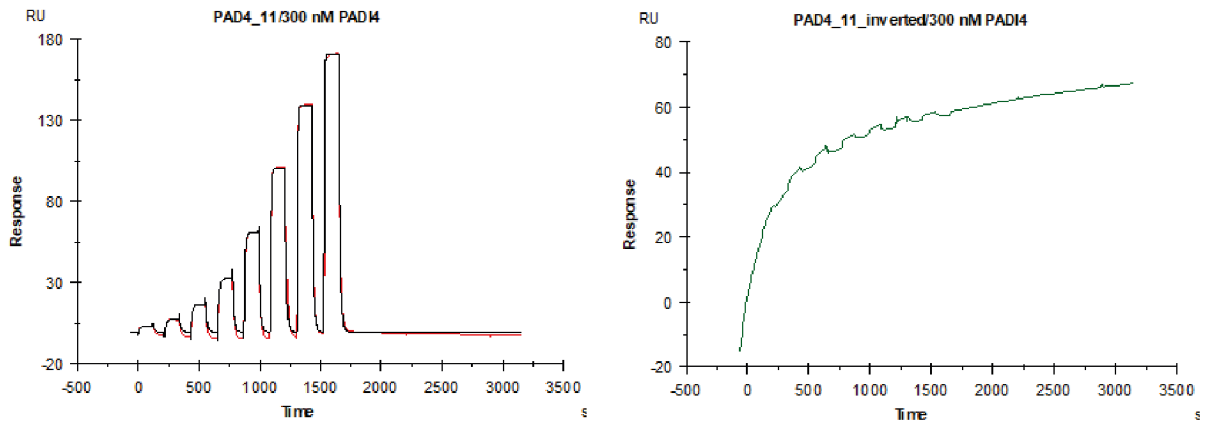

**Supplementary 10.** PADI4_11i does not bind to PADI4. SPR analysis of hPADI4 with either PADI4_11 or PADI4_11i peptides. Experiments were performed in triplicate and a representative experiment is shown.

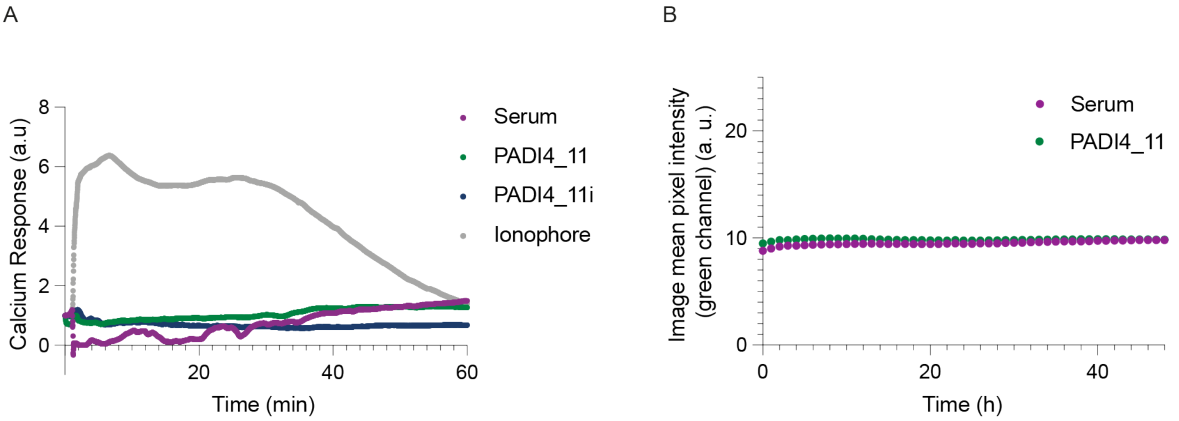

**Supplementary 11. Time course of calcium influx. A.** Calcium influx into cells over the course of 60 minutes, as measured by Calbryte intensity, after treatment with 25 µM PADI4_11 or PADI4_11i. Calcium ionophore (10 µM) used as a positive control for calcium influx. **B.** Assessment of cytotoxicity by live cell imaging with Incucyte® Cytotox Green Dye, as a read-out for cell death. hPADI4 expressing mES cells treated with 25µM PADI4_11 or DMSO vehicle (0.1%), and imaged for 48 h.

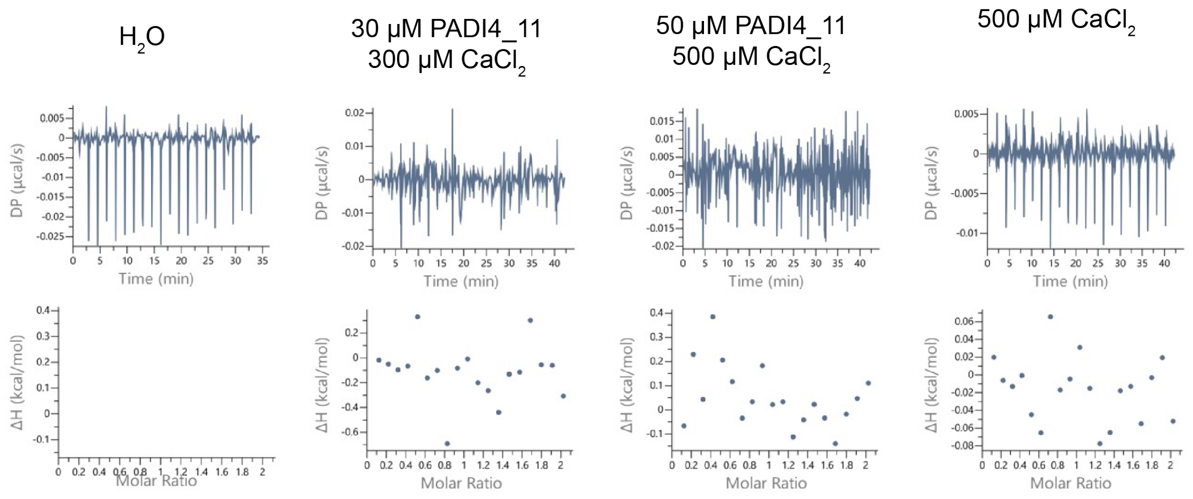

**Supplementary Figure 12. PADI4_11 does not bind to calcium.** Isothermal Titration Calorimetry between PADI4_11 peptide (30 and 50 µM) and $CaCl_2$ (300 and 500 µM).

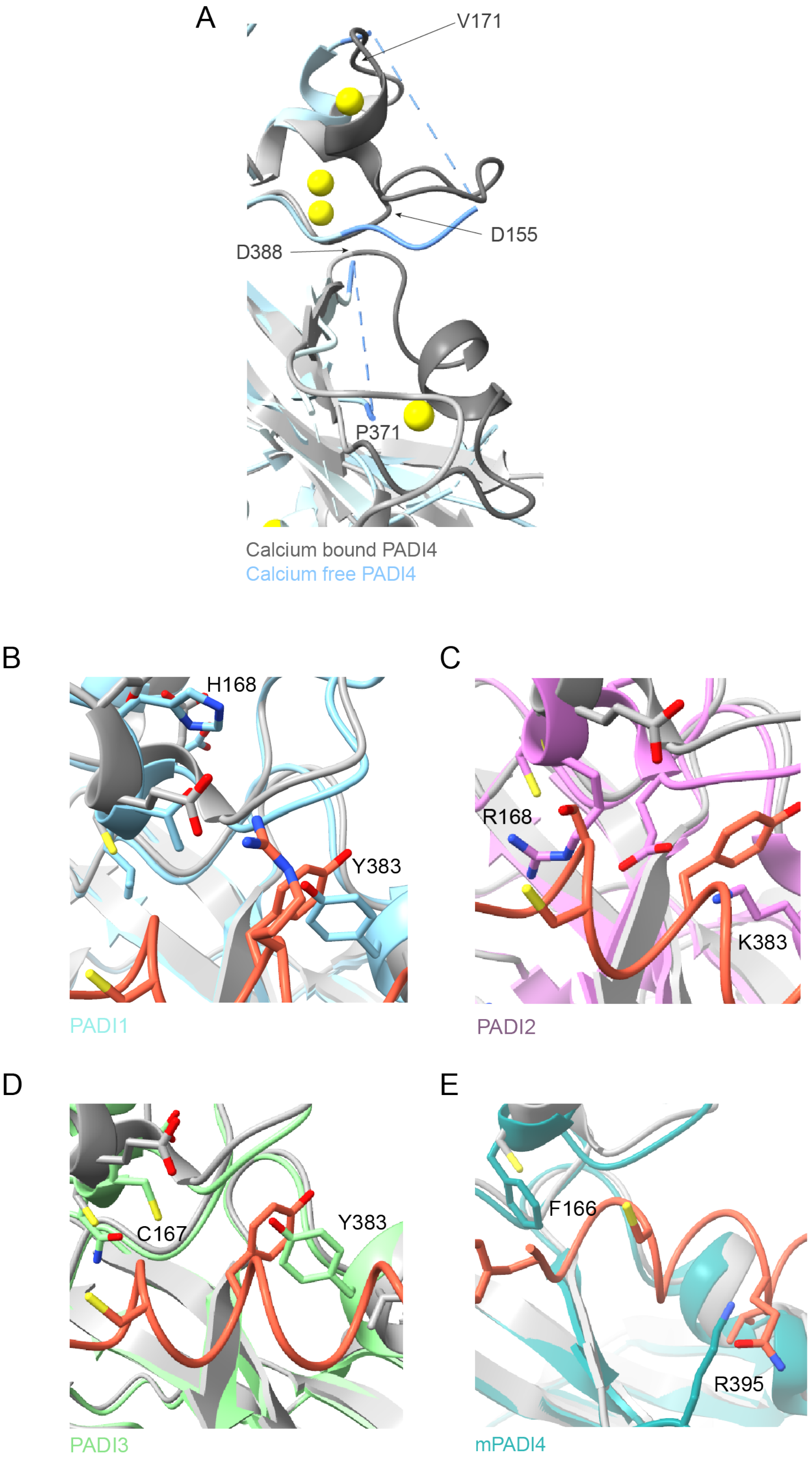

**Supplementary Figure 13. PADI4_11 binds selectively to hPADI4. A.** Alignment of PADI4 bound to calcium (grey) and calcium free PADI4 ( PBD ID: 1WD8) (blue). Loops involved in the binding of PADI4_11 peptide are shown in dark grey. **B.** Structure alignment of PADI1 AlphaFold model (cyan) with hPADI4 in complex with peptide PADI4_11. **C.** Structure alignment of PADI2 (PDB: 4N2B, magenta) with hPADI4 in complex with peptide PADI4_11.

**D.** Structure alignment of PADI3 AlphaFold model (light green) with hPADI4 in complex with peptide PADI4_11. **E.** Structure alignment of mouse PADI4 AlphaFold model (teal) with hPADI4 in complex with peptide PADI4_11. Protein residues predicted to clash with PADI4_11 are highlighted as sticks.

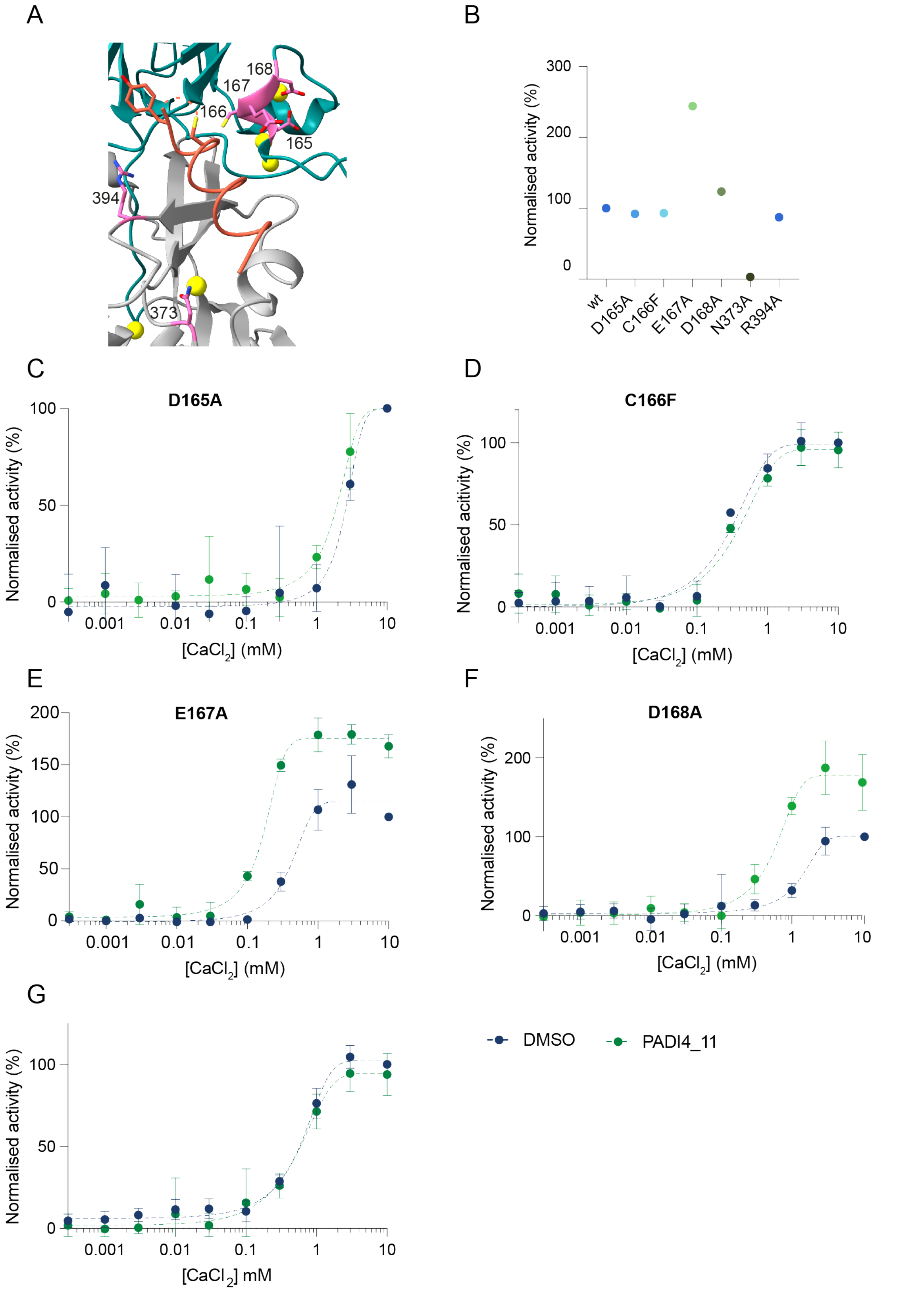

**Supplementary Figure 14. PADI4_11 cannot activate some hPADI4 variants. A**. View from a PADI4 cryoEM structure with PADI4_11 (orange). Residues involved in the binding that were mutated for following experiments are shown in pink. **B.** Different PADI4 variants have different activities. Activity of each PADI4 variant was measured by COLDER assay at 10 mM $CaCl_2$ following 30 min incubation at RT. Activity was normalised against the activity of wild-type PADI4. **C-G**. PADI4_11 is not able to activate some PADI4 variants. COLDER assays were performed in presence or absence of PADI4_11 at 30 µM and different concentrations of $CaCl_2$. Activity was normalised against the activity of each variant at 10 mM $CaCl_2$ in absence of peptide. Data represents mean ± SEM of two independent replicates. Each replicate was performed in triplicate.

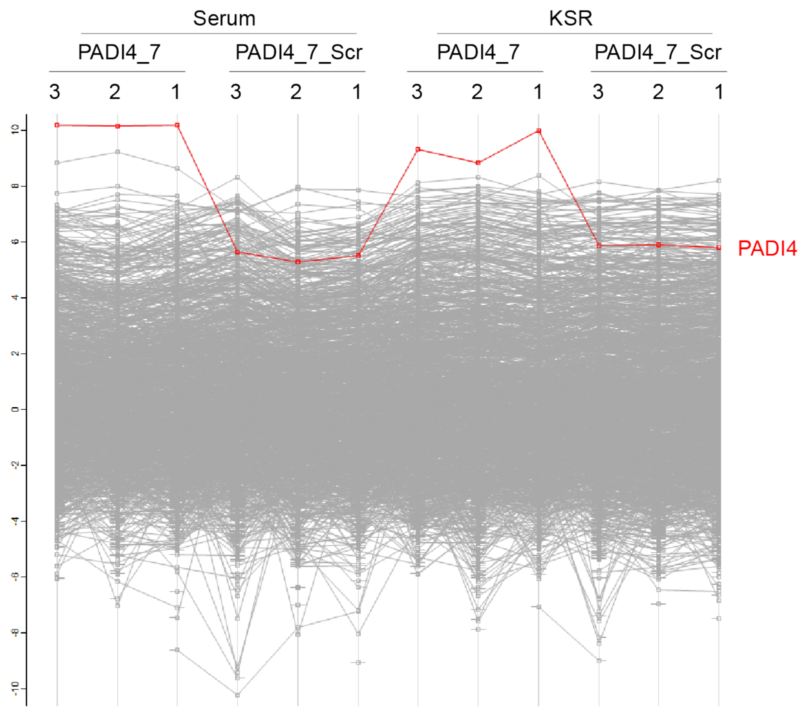

**Supplementary Figure 15. Specific enrichment of PADI4 by bio-PADI4_7.**

Enrichment profile for PADI4 compared to all other identified proteins across all pull-down samples, as determined by Mass Spectrometric analysis.

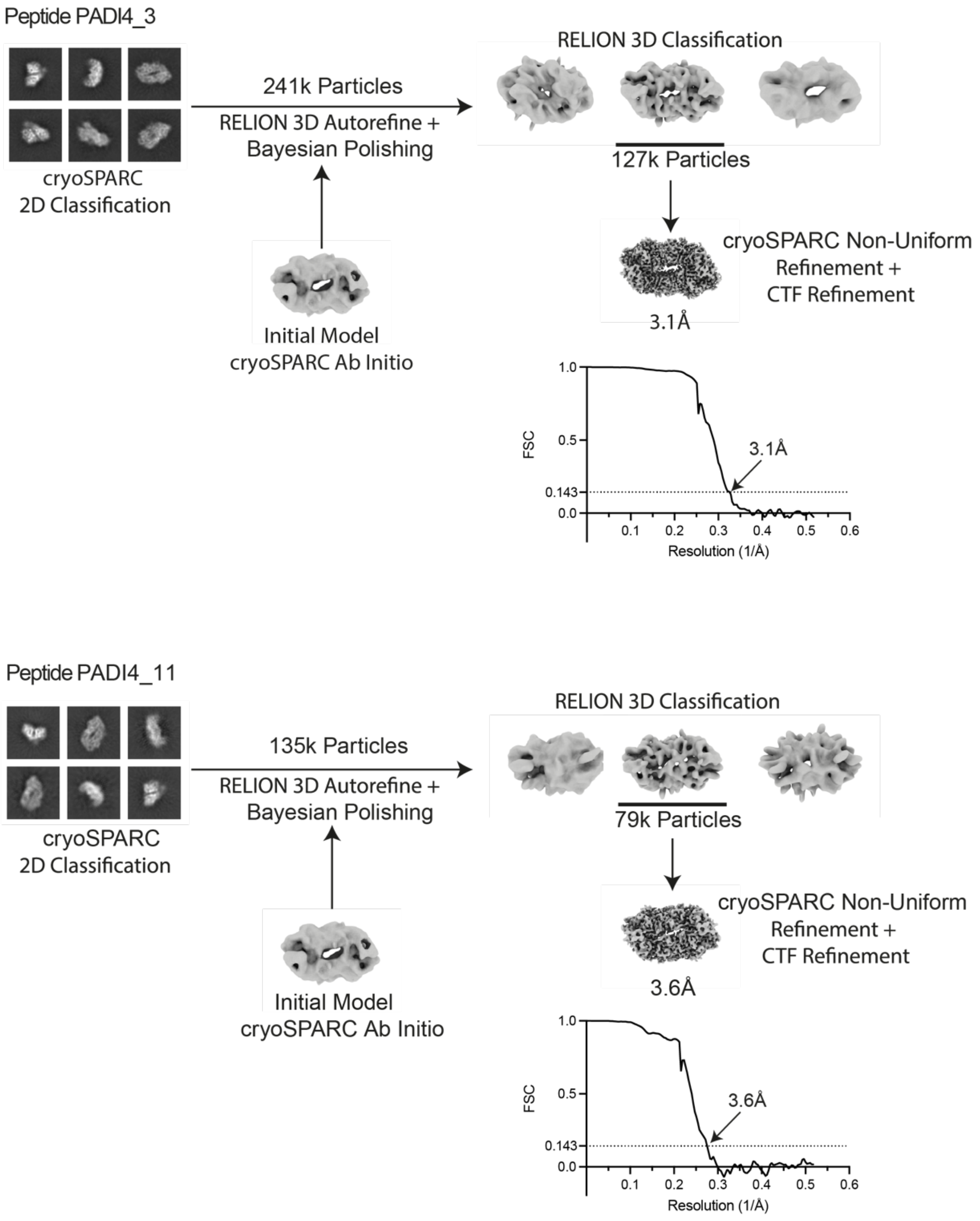

**Supplementary Figure 16.** Cryo-electron microscopy data processing pipeline for the structures of PADI4 in complex with PADI4_3 (top) and PADI4 in complex with PADI4_11 (bottom).

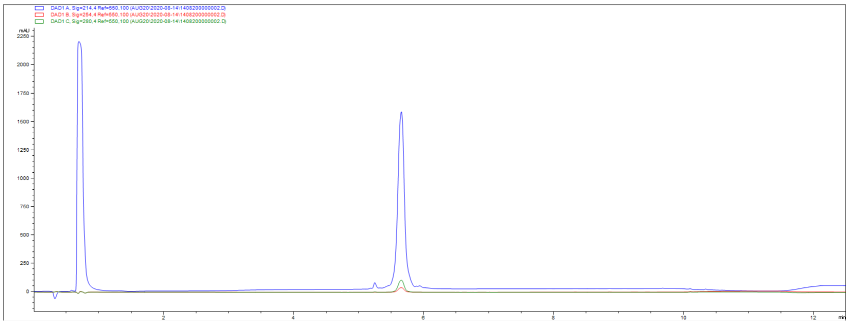

**Supplementary Figure 17**. PADI4_2 HPLC trace, showing A214 in blue, A254 in red and A280 in green.

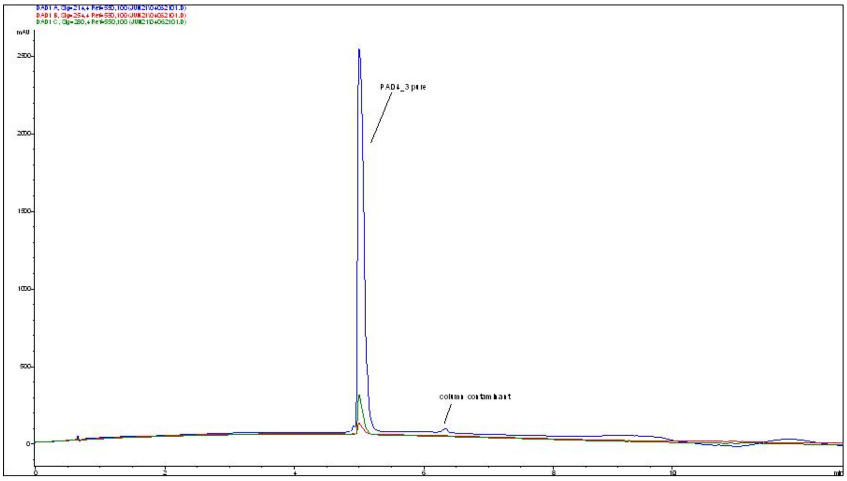

**Supplementary Figure 18**. PADI4_3 HPLC trace, showing A214 in blue, A254 in red and A280 in green.

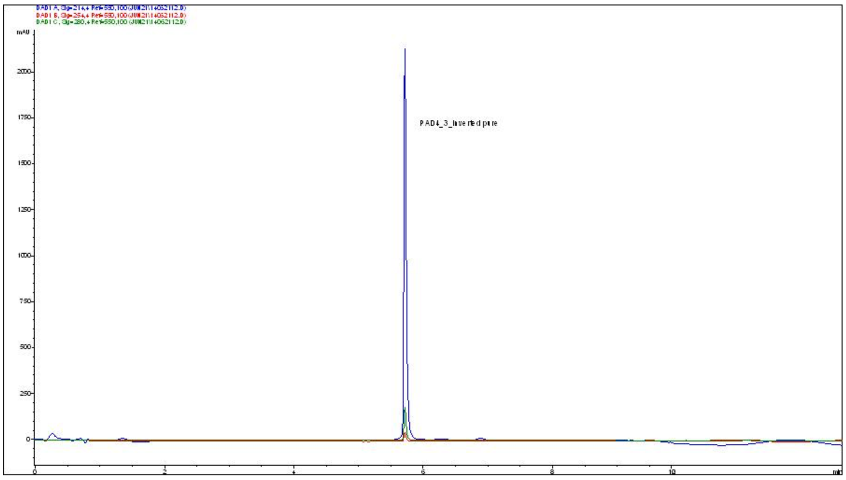

**Supplementary Figure 19**. PADI4_3i HPLC trace, showing A214 in blue, A254 in red and A280 in green.

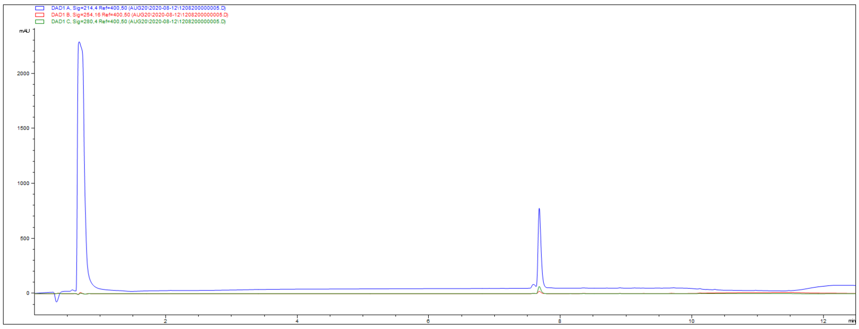

**Supplementary Figure 20**. PADI4_4 HPLC trace, showing A214 in blue, A254 in red and A280 in green.

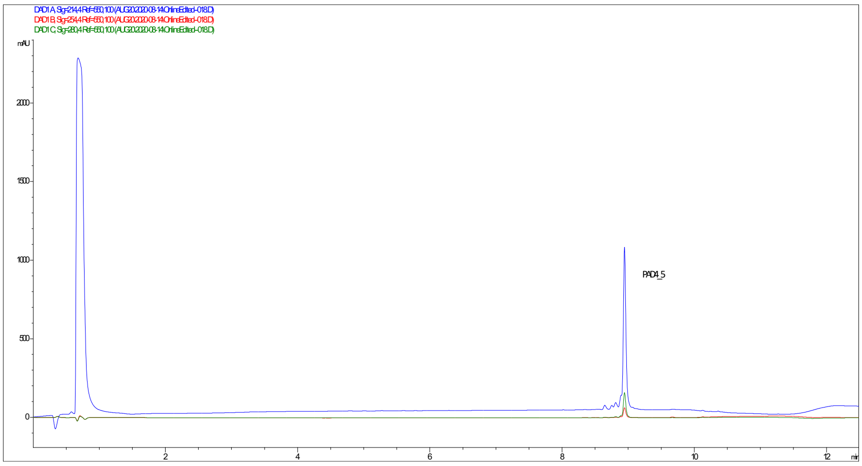

**Supplementary Figure 21**. PADI4_5 HPLC trace, showing A214 in blue, A254 in red and A280 in green.

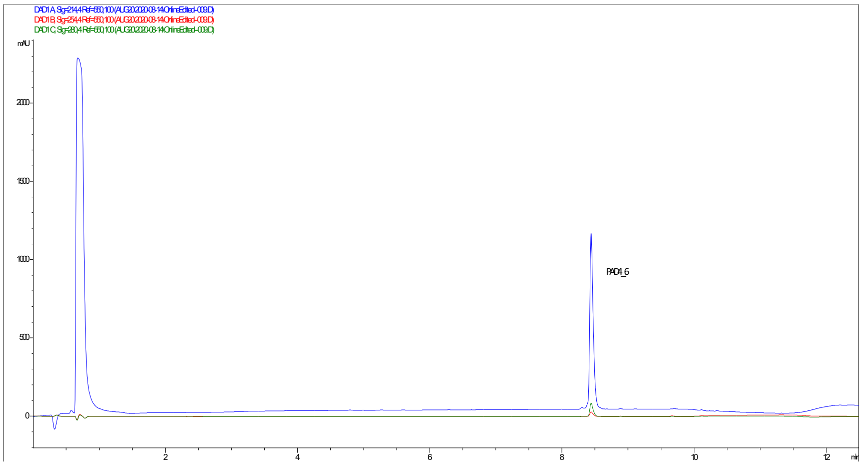

**Supplementary Figure 22**. PADI4_6 HPLC trace, showing A214 in blue, A254 in red and A280 in green.

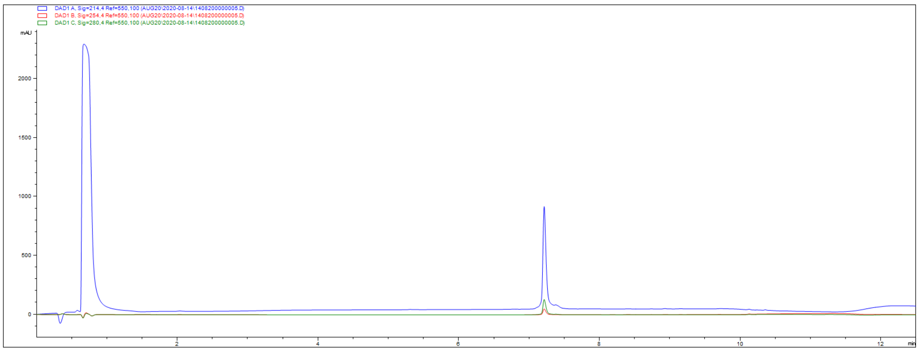

**Supplementary Figure 23**. PADI4_7 HPLC trace, showing A214 in blue, A254 in red and A280 in green.

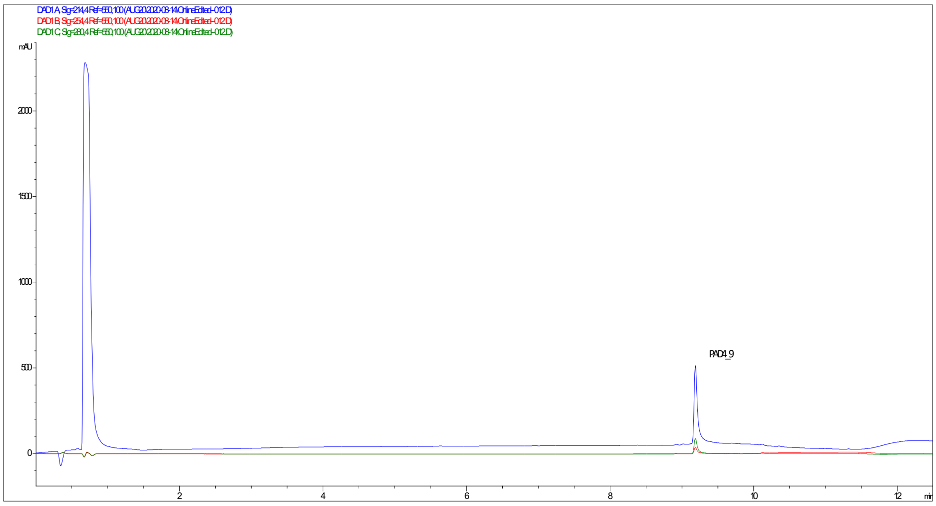

**Supplementary Figure 24**. PADI4_9 HPLC trace, showing A214 in blue, A254 in red and A280 in green.

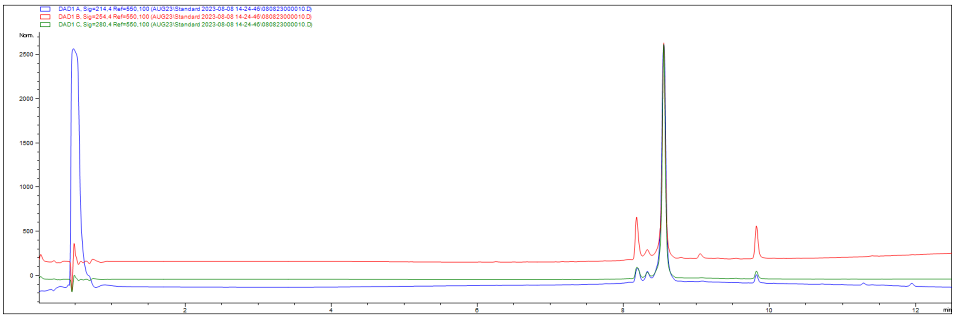

**Supplementary Figure 25**. PADI4_10 HPLC trace, showing A214 in blue, A254 in red and A280 in green.

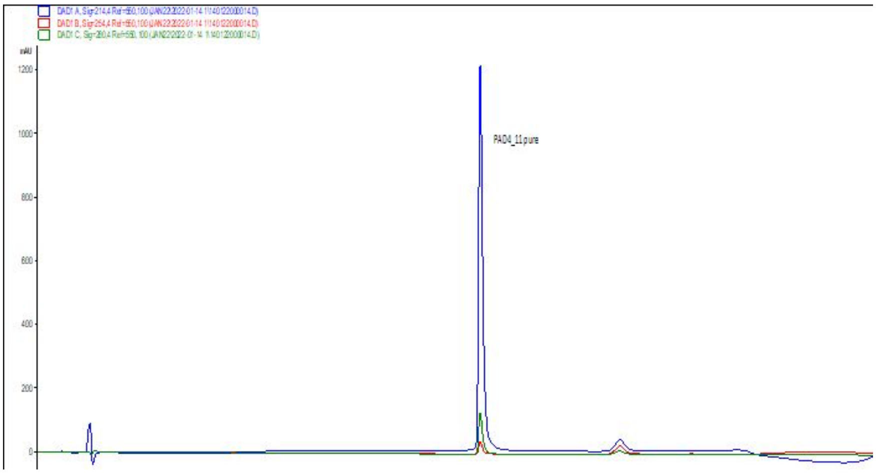

**Supplementary Figure 26**. PADI4_11 HPLC trace, showing A214 in blue, A254 in red and A280 in green.

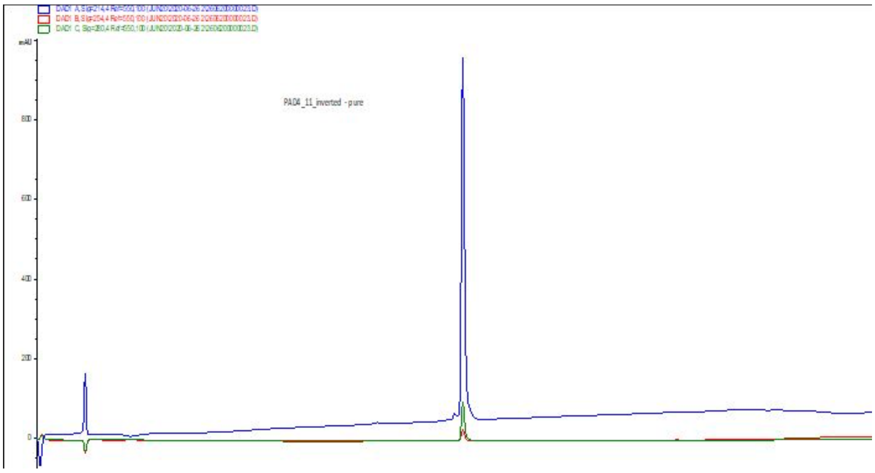

**Supplementary Figure 27**. PADI4_11i HPLC trace, showing A214 in blue, A254 in red and A280 in green.

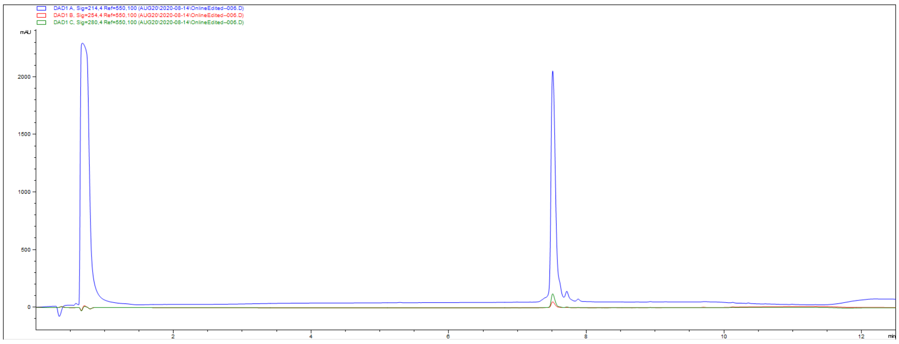

**Supplementary Figure 28**. PADI4_12 HPLC trace, showing A214 in blue, A254 in red and A280 in green.

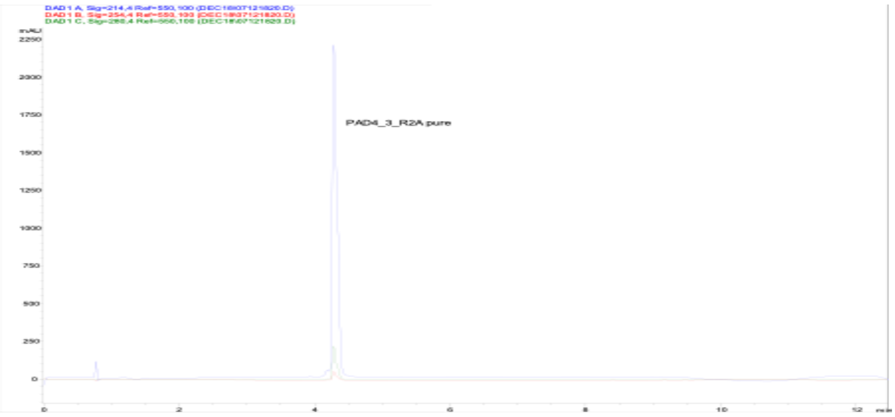

**Supplementary Figure 29**. PADI4_3_R2A HPLC trace, showing A214 in blue, A254 in red and A280 in green.

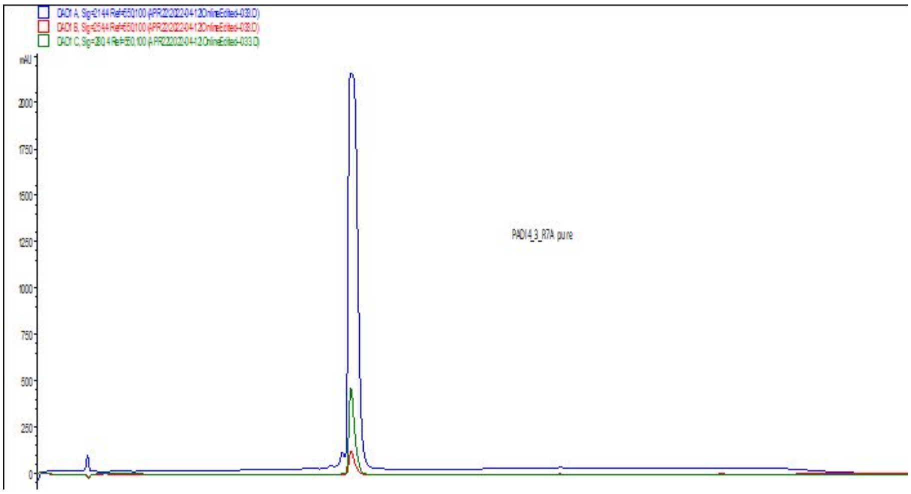

**Supplementary Figure 30**. PADI4_3_R7A HPLC trace, showing A214 in blue, A254 in red and A280 in green.

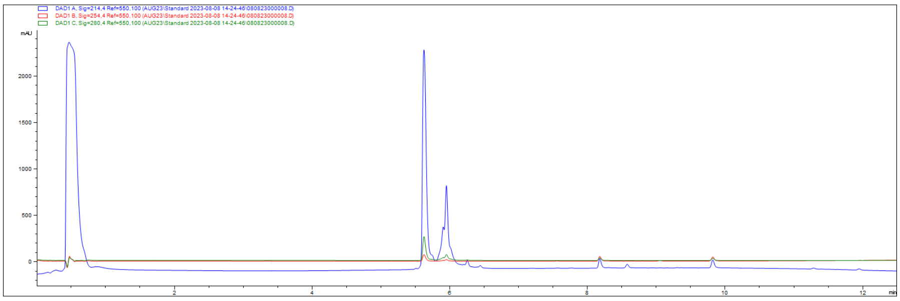

**Supplementary Figure 31**. PADI4_3_R2cit HPLC trace, showing A214 in blue, A254 in red and A280 in green.

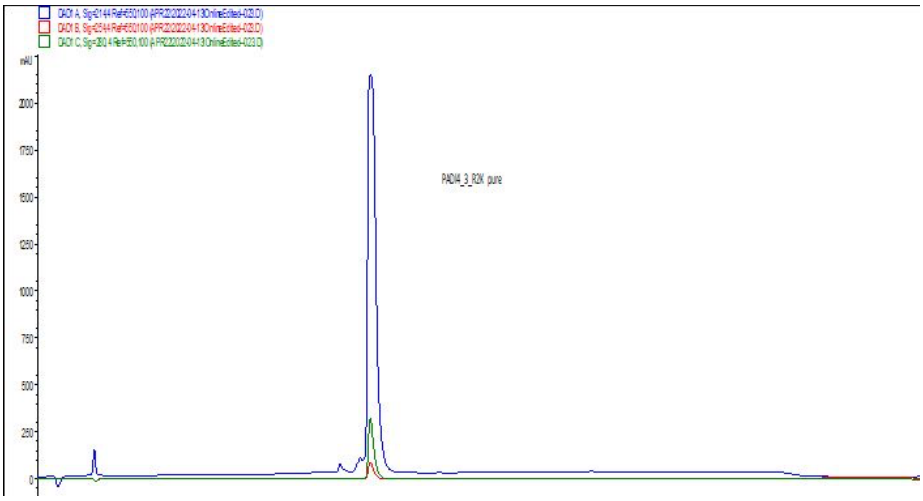

**Supplementary Figure 32**. PADI4_3_R2K HPLC trace, showing A214 in blue, A254 in red and A280 in green.

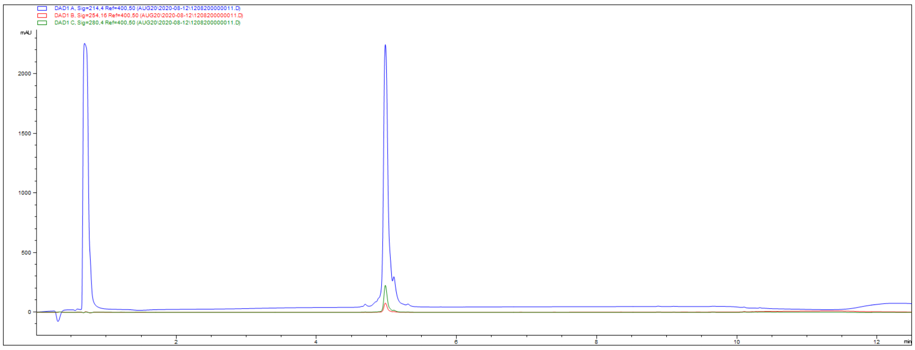

**Supplementary Figure 33**. PADI4_3_y1(OMe)Y HPLC trace, showing A214 in blue, A254 in red and A280 in green.

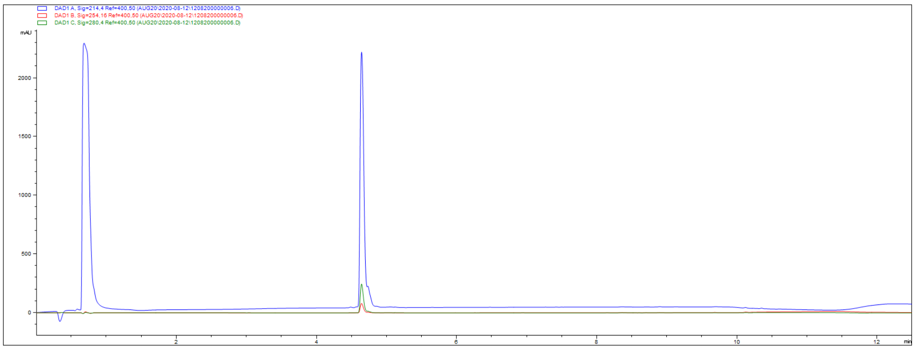

**Supplementary Figure 34**. PADI4_3_H4R HPLC trace, showing A214 in blue, A254 in red and A280 in green.

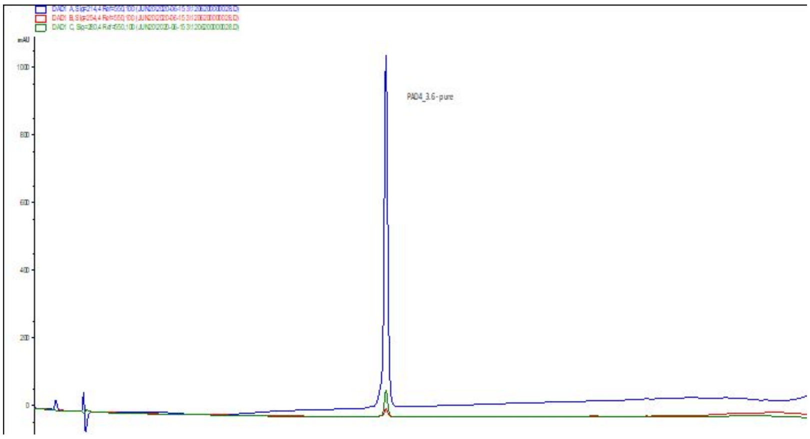

**Supplementary Figure 35**. PADI4_3_K10I HPLC trace, showing A214 in blue, A254 in red and A280 in green.

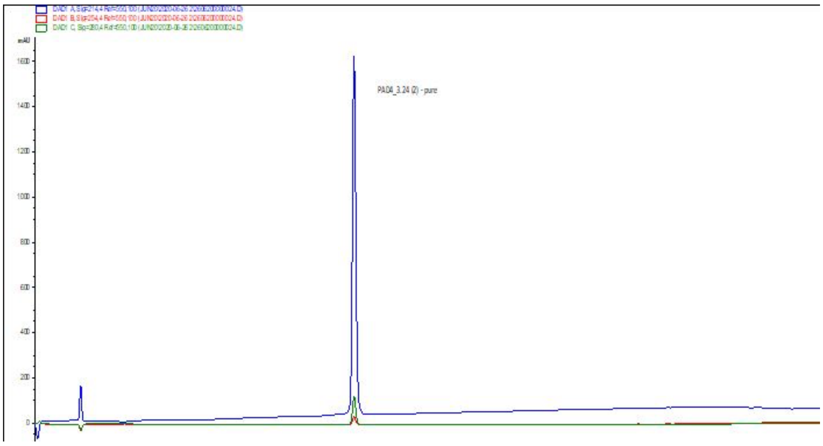

**Supplementary Figure 36**. PADI4_3_K10V HPLC trace, showing A214 in blue, A254 in red and A280 in green.

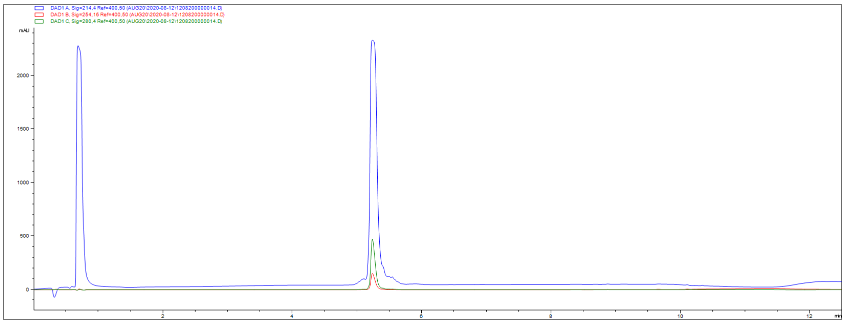

**Supplementary Figure 37**. PADI4_3_H4R_K10I HPLC trace, showing A214 in blue, A254 in red and A280 in green.

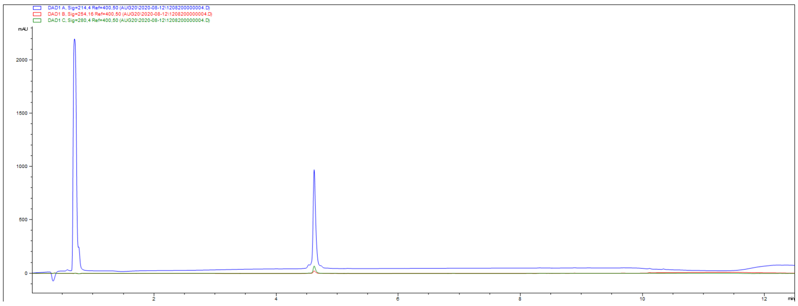

**Supplementary Figure 38**. PADI4_3_R7N HPLC trace, showing A214 in blue, A254 in red and A280 in green.

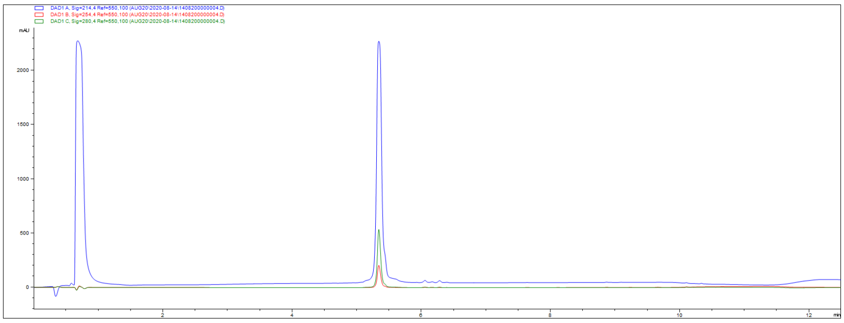

**Supplementary Figure 39**. PADI4_3_R7W HPLC trace, showing A214 in blue, A254 in red and A280 in green.

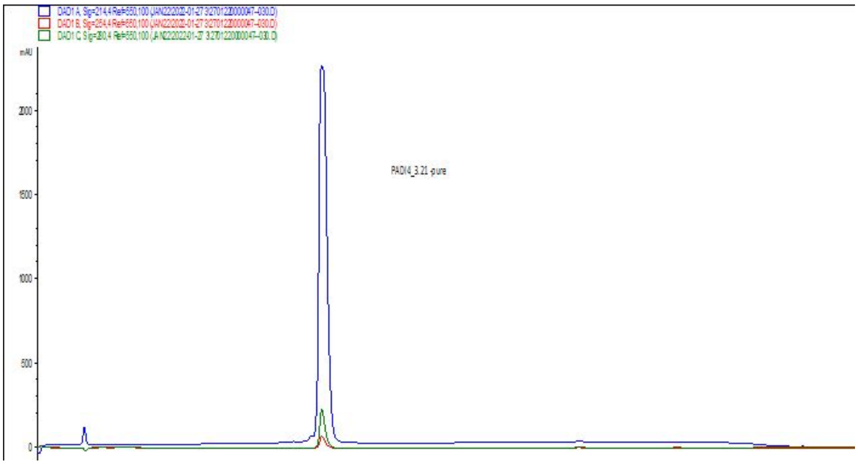

**Supplementary Figure 40**. PADI4_3_Y6H HPLC trace, showing A214 in blue, A254 in red and A280 in green.

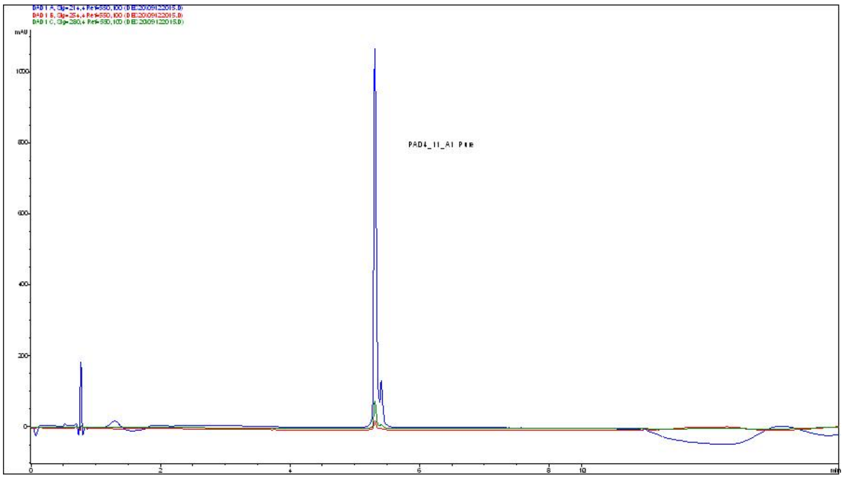

**Supplementary Figure 41**. PADI4_11_Y1A HPLC trace, showing A214 in blue, A254 in red and A280 in green.

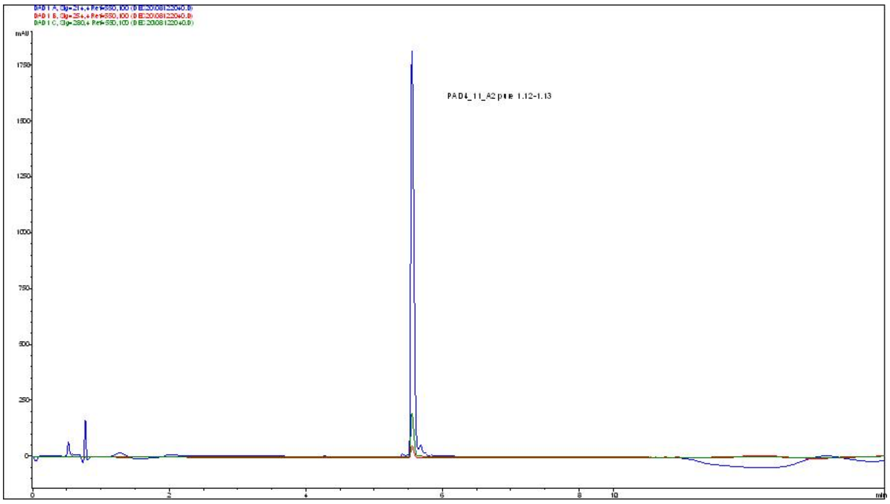

**Supplementary Figure 42**. PADI4_11_E2A HPLC trace, showing A214 in blue, A254 in red and A280 in green.

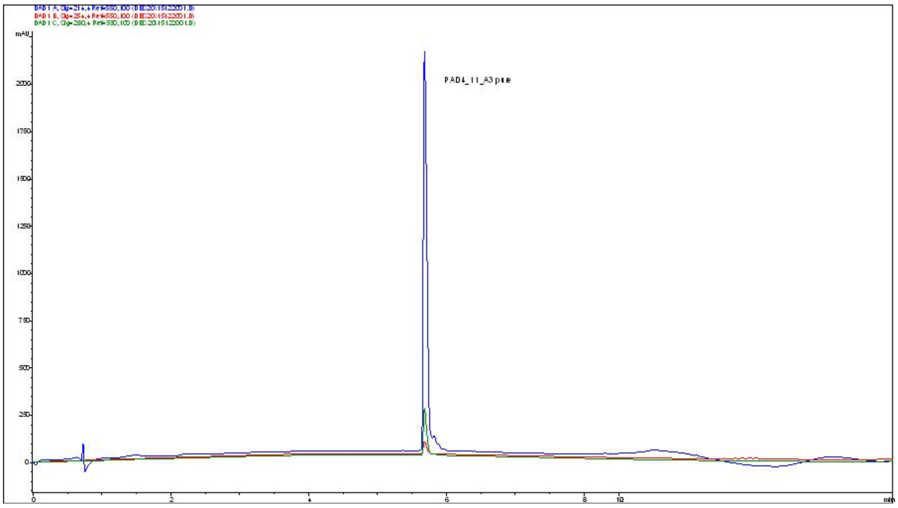

**Supplementary Figure 43**. PADI4_11_S3A HPLC trace, showing A214 in blue, A254 in red and A280 in green.

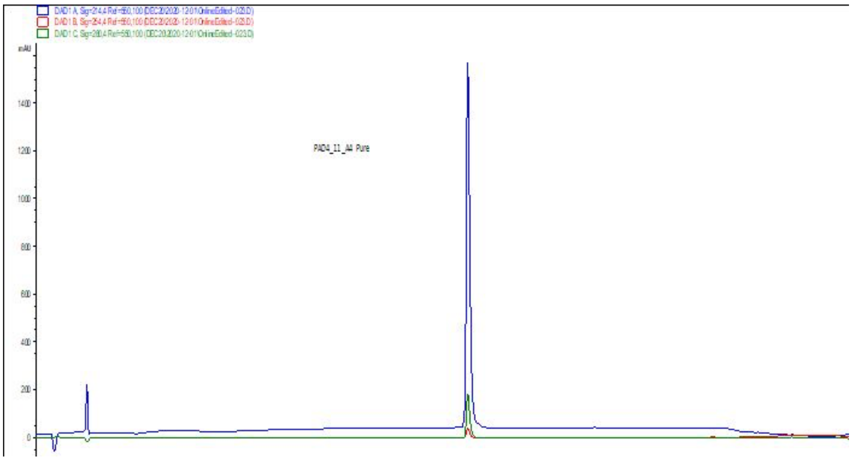

**Supplementary Figure 44**. PADI4_11_C4A HPLC trace, showing A214 in blue, A254 in red and A280 in green.

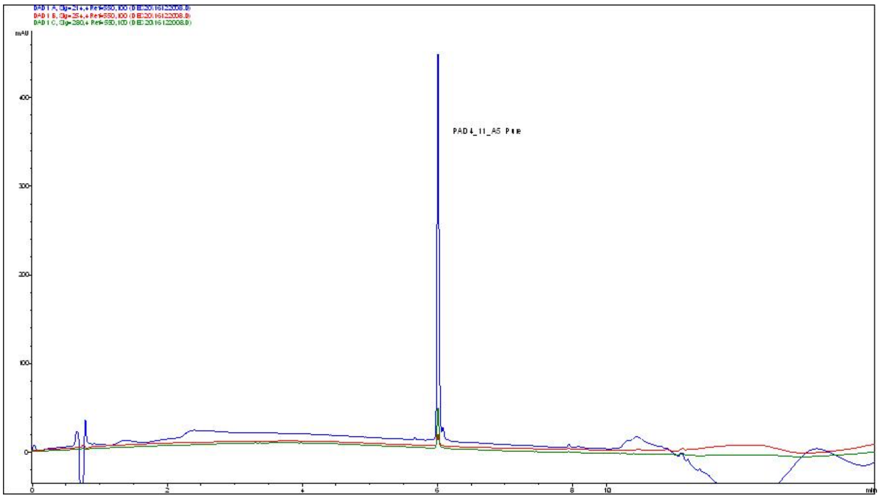

**Supplementary Figure 45**. PADI4_11_R5A HPLC trace, showing A214 in blue, A254 in red and A280 in green.

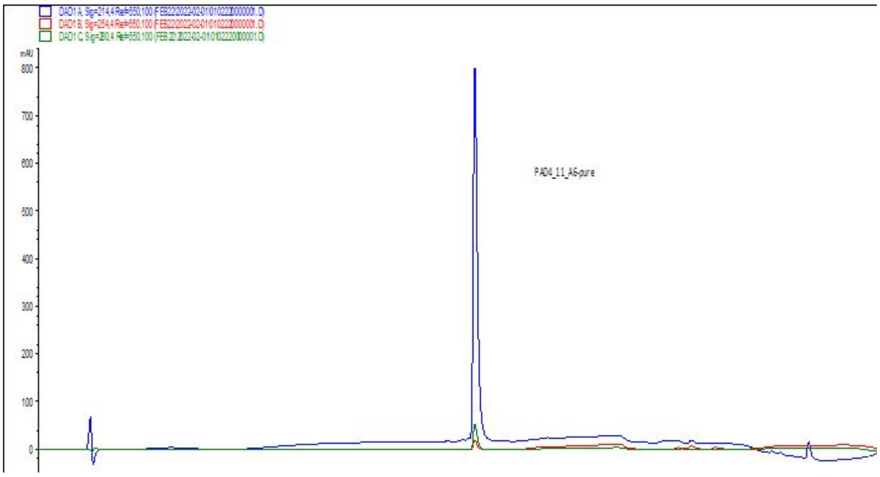

**Supplementary Figure 46**. PADI4_11_Y6A HPLC trace, showing A214 in blue, A254 in red and A280 in green.

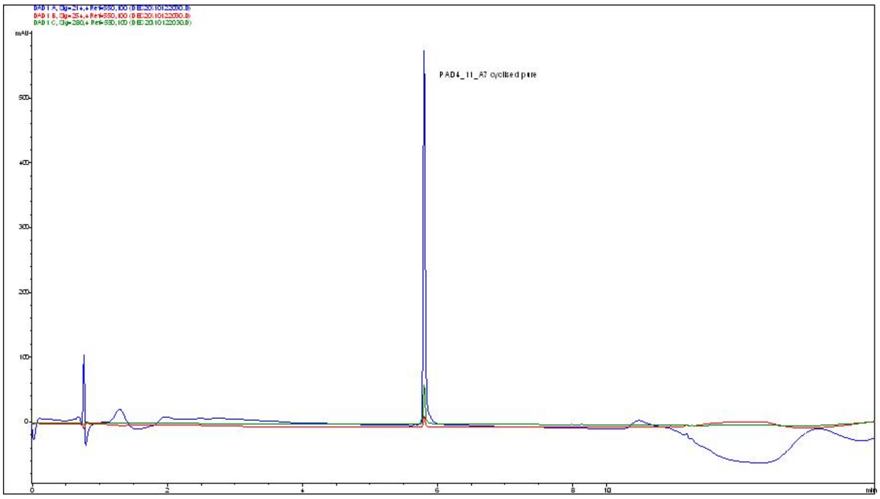

**Supplementary Figure 47**. PADI4_11_R7A HPLC trace, showing A214 in blue, A254 in red and A280 in green.

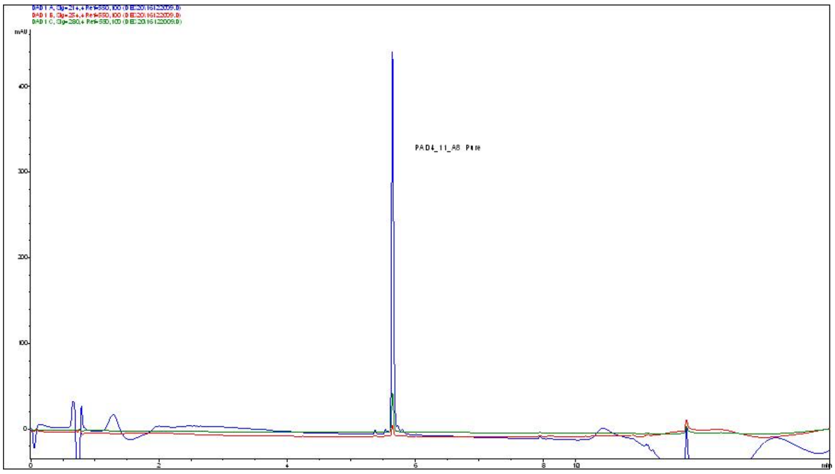

**Supplementary Figure 48**. PADI4_11_Q8A HPLC trace, showing A214 in blue, A254 in red and A280 in green.

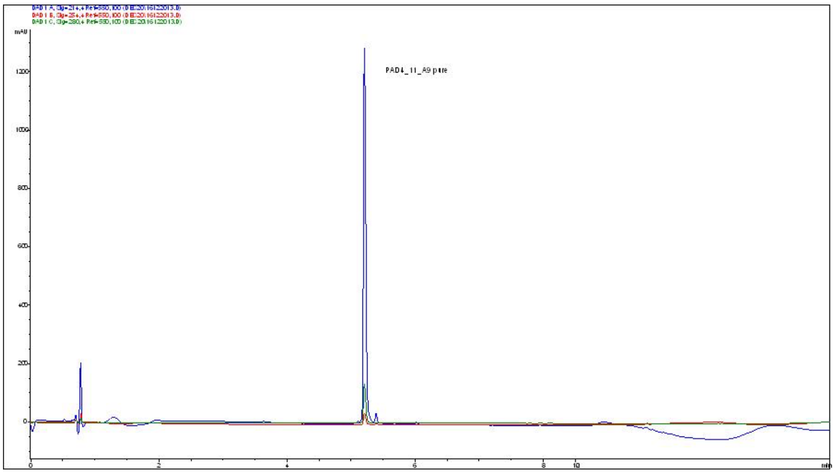

**Supplementary Figure 49**. PADI4_11_V9A HPLC trace, showing A214 in blue, A254 in red and A280 in green.

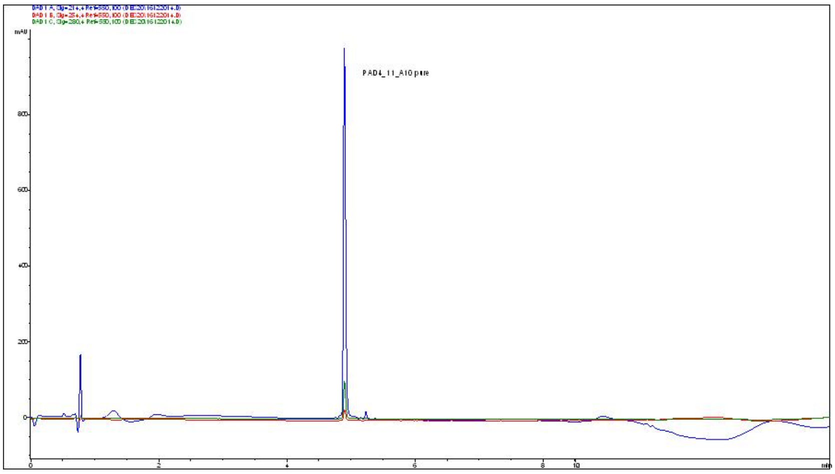

**Supplementary Figure 50.** PADI4_11_L10A HPLC trace, showing A214 in blue, A254 in red and A280 in green

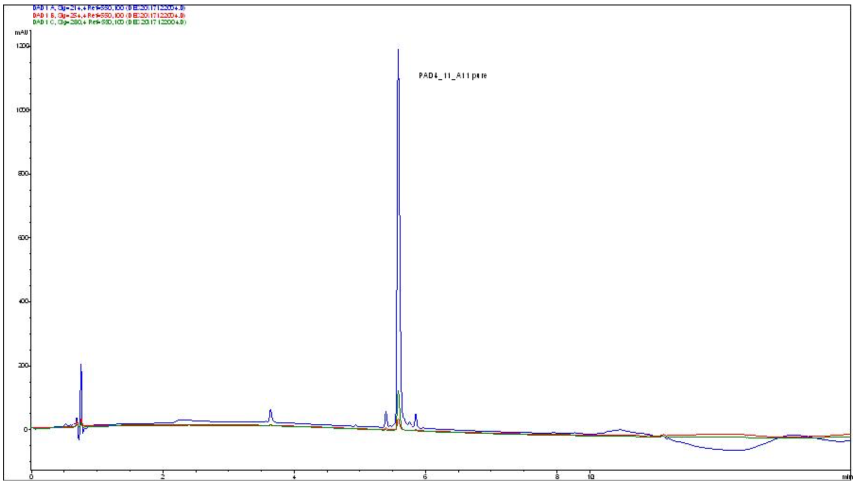

**Supplementary Figure 51**. PADI4_11_Q11A HPLC trace, showing A214 in blue, A254 in red and A280 in green.

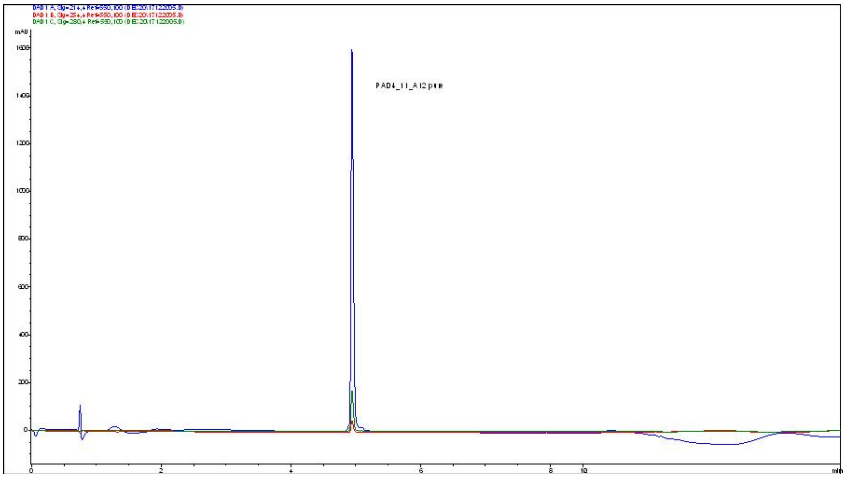

**Supplementary Figure 52**. PADI4_11_L12A HPLC trace, showing A214 in blue, A254 in red and A280 in green.

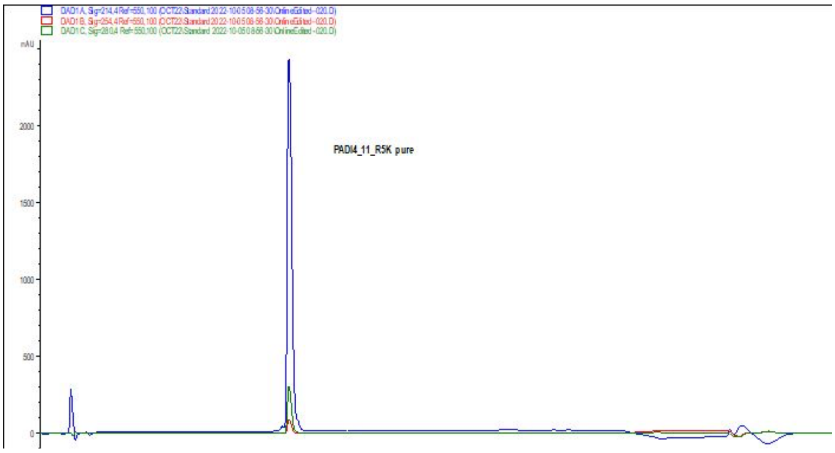

**Supplementary Figure 53**. PADI4_11_R5K HPLC trace, showing A214 in blue, A254 in red and A280 in green.

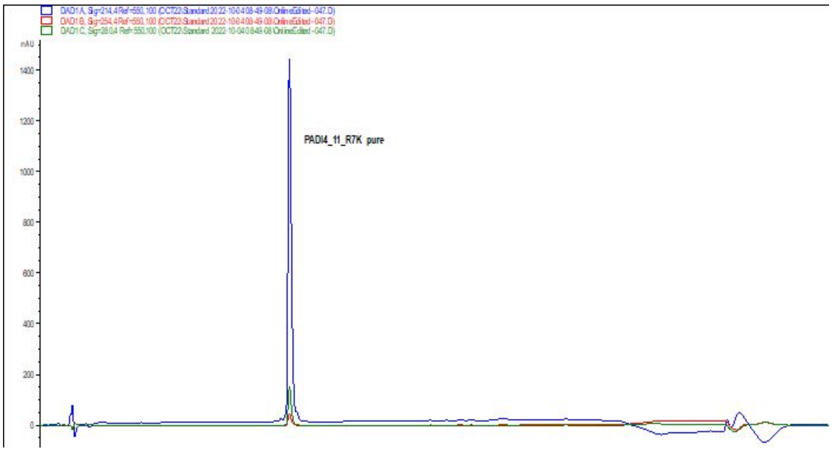

**Supplementary Figure 54**. PADI4_11_R7K HPLC trace, showing A214 in blue, A254 in red and A280 in green.

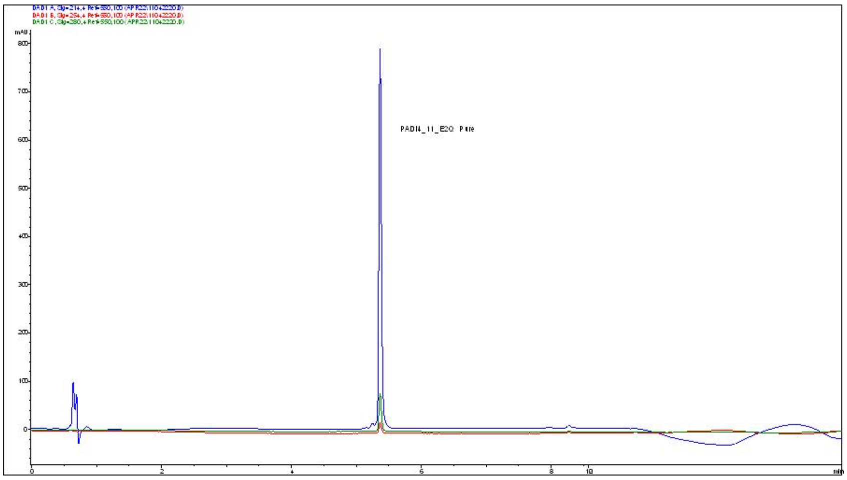

**Supplementary Figure 55**. PADI4_11_E2Q HPLC trace, showing A214 in blue, A254 in red and A280 in green.

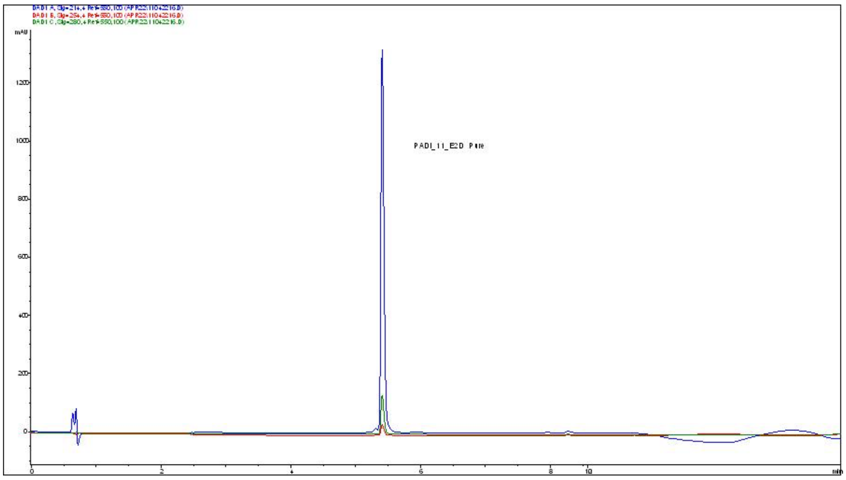

**Supplementary Figure 56**. PADI4_11_E2D HPLC trace, showing A214 in blue, A254 in red and A280 in green.

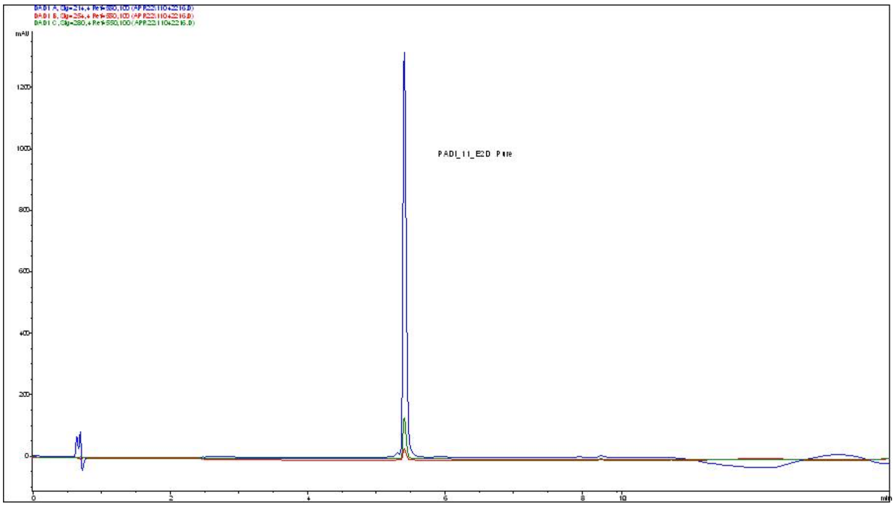

**Supplementary Figure 57**. PADI4_11B HPLC trace, showing A214 in blue, A254 in red and A280 in green.

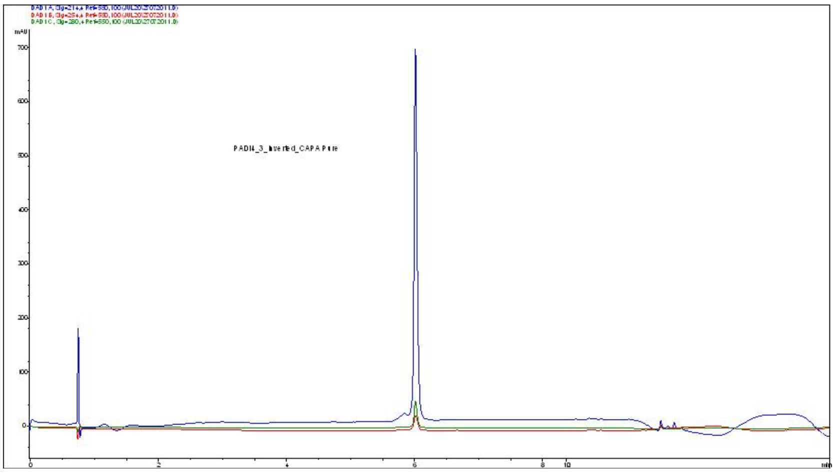

**Supplementary Figure 58**. PADI4_3_CAPA HPLC trace, showing A214 in blue, A254 in red and A280 in green.

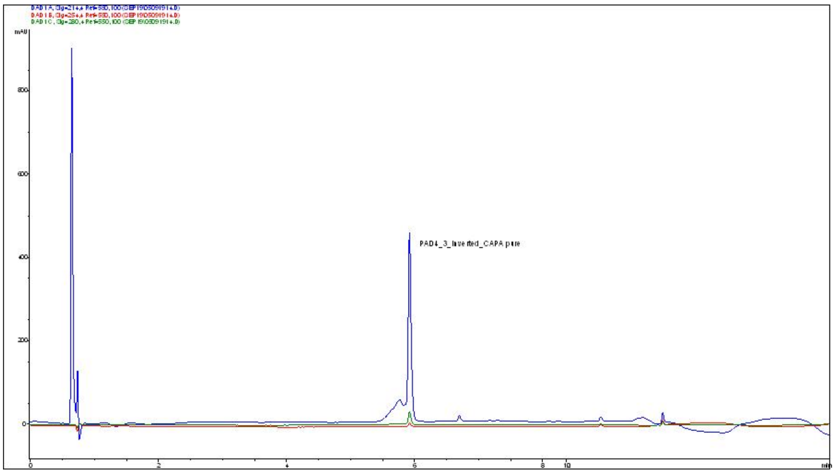

**Supplementary Figure 59**. PADI4_3i_CAPA HPLC trace, showing A214 in blue, A254 in red and A280 in green.

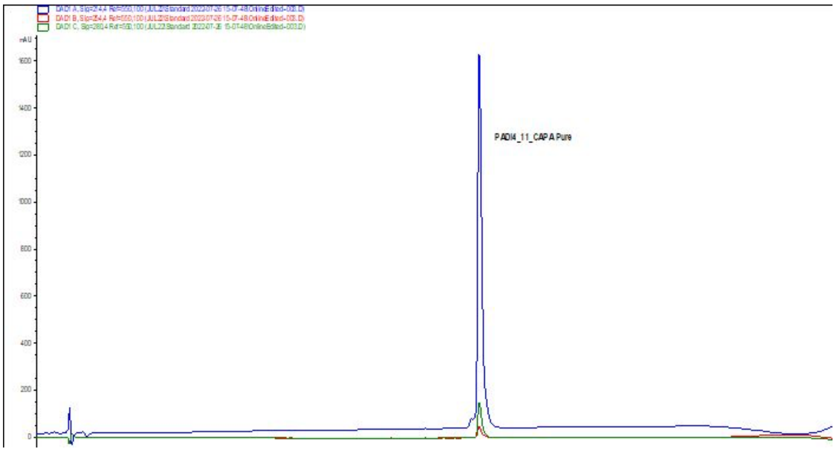

**Supplementary Figure 60**. PADI4_11_CAPA HPLC trace, showing A214 in blue, A254 in red and A280 in green

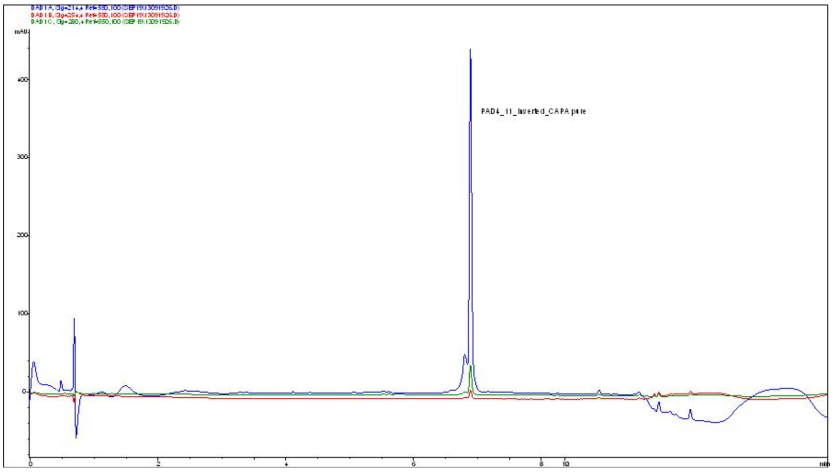

**Supplementary Figure 61**. PADI4_11i_CAPA HPLC trace, showing A214 in blue, A254 in red and A280 in green

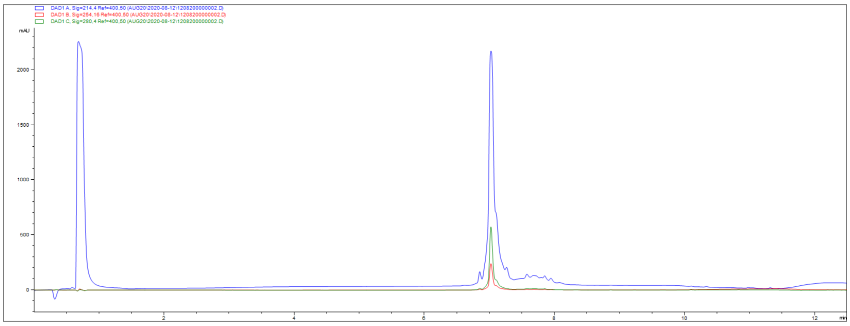

**Supplementary Figure 62**. PADI4_7_bio HPLC trace, showing A214 in blue, A254 in red and A280 in green.

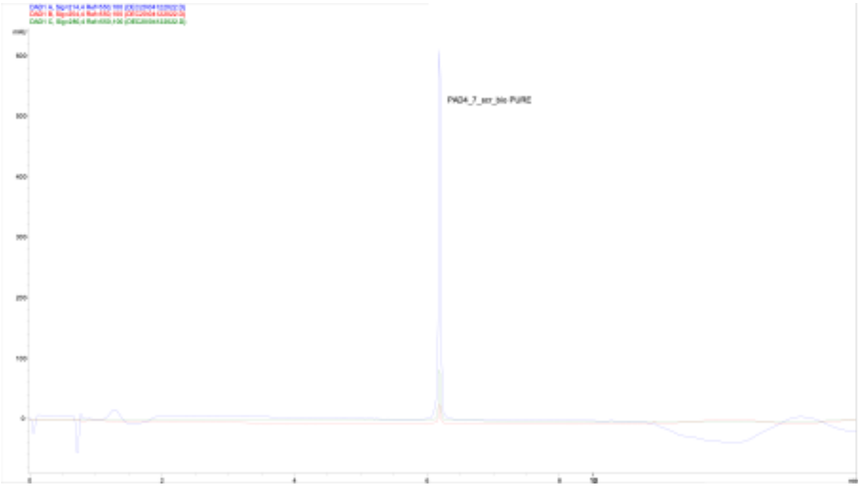

**Supplementary Figure 63**. PADI4_7scr_bio HPLC trace, showing A214 in blue, A254 in red and A280 in green.

**Supplementary Table 1. SPR data of selected macrocyclic peptides enriched in the RaPID selections.** Data shows average ± standard deviation of three independent experiments.

| | 10 mM $Ca^{2+}$ | | | 0 mM $Ca^{2+}$ | | | 200 µM Cl-amidine + 10 mM $Ca^{2+}$ | | |
|---|---|---|---|---|---|---|---|---|---|
| | Kd (nM) | Kon (1/Ms) | Koff (1/s) | Kd (nM) | Kon (1/Ms) | Koff (1/s) | Kd (nM) | Kon (1/Ms) | Koff (1/s) |
| **PAD4_1** | 205 ± 52 | $7.6\cdot10^{4}$ ± $0.8\cdot.10^{4}$ | $2\cdot10^{-2}$ ± $3\cdot10^{-3}$ | 105 ± 17 | $9.5\cdot10^{4}$ ± $0.6\cdot10^{4}$ | $1\cdot10^{-2}$ ± $2\cdot10^{-3}$ | 728 ± 175 | $4\cdot10^{4}$ ± $3\cdot10^{3}$ | $3\cdot10^{-2}$ ± $5\cdot10^{-3}$ |
| **PAD4_2** | 13 ± 4 | $8\cdot10^{4}$ ± $2\cdot10^{4}$ | $1.0\cdot10^{-3}$ ± $0.5\cdot10^{-3}$ | >5 µM | - | - | >5 µM | - | - |
| **PAD4_3** | 2.7 ± 0.5 | $2.5\cdot10^{5}$ ± $0.2\cdot10^{5}$ | $7\cdot10^{-4}$ ± $1\cdot10^{-4}$ | >5 µM | - | - | >5 µM | - | - |
| **PAD4_7** | 39 ± 24 | $5\cdot10^{6}$ ± $5\cdot10^{6}$ | $2\cdot10^{-1}$ ± $2\cdot10^{-1}$ | 9 ± 1 | $1\cdot10^{7}$ ± $1\cdot10^{7}$ | $1\cdot10^{-1}$± $1\cdot10^{-1}$ | 36 ± 12 | $9\cdot10^{6}$ ± $6\cdot10^{6}$ | $4\cdot10^{-1}$ ± $2\cdot10^{-1}$ |
| **PAD4_11** | 457 ± 109 | $2\cdot10^{5}$ ± $7\cdot10^{5}$ | $1\cdot10^{-1}$ ± $2\cdot10^{-2}$ | >5 µM | - | - | 382 ± 27 | $3\cdot10^{5}$ ± $5\cdot10^{4}$ | $1\cdot10^{-1}$ ± $1\cdot10^{-2}$ |
| **PAD4_12** | 679 ± 83 | $2\cdot10^{5}$ ± $5\cdot10^{4}$ | $1\cdot10^{-1}$ ± $3\cdot10^{-2}$ | >5 µM | - | - | 674 ± 191 | $3\cdot10^{5}$ ± $8\cdot10^{4}$ | $2\cdot10^{-1}$ ± $2\cdot10^{-2}$ |

**Supplementary Table 2. Cryo-EM data collection, refinement, and validation statistics.**

| | PADI4_3 (EMDB-19011) (PDB 8R8U) | PADI4_11 (EMDB-19012) (PDB 8R8V) |
|---|---|---|
| **Data collection and processing** | | |
| Voltage (kV) | 300 | 300 |
| Electron exposure (e–/Å$^2$) | 28.0 | 28.0 |
| Defocus range (μm) | -1 – -3 | -1 – -3 |
| Pixel size (Å) | 0.95 | 0.95 |
| Symmetry imposed | C2 | C2 |
| Final particle images (no.) | 127k | 79k |
| Map resolution (Å) | 3.1 | 3.6 |
| FSC threshold 0.143 | | |
| **Refinement** | | |
| Initial model used (PDB code) | 1WD9 | 1WD9 |
| Model resolution (Å) | 3.2 | 3.7 |
| FSC threshold 0.5 | | |
| Map sharpening *B* factor (Å$^2$) | -146.3 | -187.6 |
| Model composition | | |
| Non-hydrogen atoms | 10140 | 10198 |
| Protein residues | 1282 | 1292 |
| Ligands | 10 | 10 |
| *B* factors (Å$^2$) | | |
| Protein | 46.7 | 63.1 |
| Ligand | 29.6 | 49.2 |
| R.m.s. deviations | | |
| Bond lengths (Å) | 0.002 | 0.005 |
| Bond angles (°) | 0.456 | 0.708 |
| Validation | | |
| MolProbity score | 1.20 | 1.64 |
| Clashscore | 4.12 | 7.29 |
| Poor rotamers (%) | 0.18 | 0.26 |
| Ramachandran plot | | |
| Favored (%) | 98.01 | 96.38 |
| Allowed (%) | 1.99 | 3.62 |
| Disallowed (%) | 0.00 | 0.00 |

**Supplementary Table 3. SPR binding affinities for the different PADI4_3 analogues**. Data represents average ± standard deviation of three independent replicates.

| | Sequence | $K_D$ (nM) | Ka (1/Ms) | Kd (1/s) |
|---|---|---|---|---|
| **PADI4_3_Y1Y(Ome)** | dY(OMe)RDHHYRHPKYCG | 23 ± 2 | $1 \cdot 10^{05} \pm 1 \cdot 10^{04}$ | $3 \cdot 10^{-03} \pm 2 \cdot 10^{-05}$ |
| **PAD4_3_R2A** | dYADHHYRHPKYCG | 239 ± 8 | $2 \cdot 10^{05} \pm 5 \cdot 10^{04}$ | $3 \cdot 10^{-02} \pm 2 \cdot 10\text{-}^{02}$ |
| **PAD4_3_R2cit** | dYcitDHHYRHPKYCG | 20 ± 11 | $5 \cdot 10^{05} \pm 6 \cdot 10^{05}$ | $2 \cdot 10^{-03} \pm 2 \cdot 10^{-04}$ |
| **PADI4_3_R2K** | dYKDHHYRHPKYCG | 9 ± 2 | $2 \cdot 10^{05} \pm 2 \cdot 10^{04}$ | $2 \cdot 10^{-03} \pm 4 \cdot 10^{-04}$ |
| **PADI4_3_H4R** | dYRDRHYRHPKYCG | 14 ± 6 | $1 \cdot 10^{06} \pm 1 \cdot 10^{06}$ | $1 \cdot 10^{-02} \pm 7 \cdot 10^{-03}$ |
| **PADI4_3_Y6H** | dYRDHHHRHPKYCG | 30 ± 15 | $6 \cdot 10^{04} \pm 3 \cdot 10^{04}$ | $1 \cdot 10^{-03} \pm 2 \cdot 10\text{-}_{04}$ |
| **PADI4_3_R7N** | dYRDHHYNHPKYCG | 41 ± 18 | $1 \cdot 10^{04} \pm 6 \cdot 10^{03}$ | $4 \cdot 10^{-04} \pm 1 \cdot 10^{-05}$ |
| **PADI4_3_R7W** | dYRDHHYWHPKYCG | 6 ± 3 | $2 \cdot 10^{05} \pm 1 \cdot 10^{05}$ | $2 \cdot 10^{-03} \pm 2 \cdot 10^{-03}$ |
| **PAD4_3_R7A** | dYRDHHYAHPKYCG | 7 ± 1 | $7 \cdot 10^{04} \pm 2 \cdot 10^{04}$ | $5 \cdot 10^{-04} \pm 7 \cdot 10^{-05}$ |
| **PADI4_3_K10I** | dYRDHHYRHPIYCG | 4 ± 2 | $1 \cdot 10^{05} \pm 4 \cdot 10^{04}$ | $4 \cdot 10^{-04} \pm 2 \cdot 10^{-04}$ |
| **PADI4_3_K10V** | dYRDHHYRHPVYCG | 5 ± 2 | $8 \cdot 10^{04} \pm 2 \cdot 10^{04}$ | $4 \cdot 10^{-04} \pm 4 \cdot 10^{-05}$ |
| **PADI4_3H4R_K10_I** | dYRDRHYRHPIYCG | 7 ± 3 | $7 \cdot 10^{05} \pm 4 \cdot 10^{05}$ | $2 \cdot 10^{-03} \pm 2 \cdot 10^{-03}$ |

**Supplementary Table 4. Primers used in this study.** SP18 is an 18-atom hexa-ethyleneglycol spacer.

| Primer Name | Sequence |
|---|---|
| *mPadi4*_fw | CATATGATGGCCCAGGGTGCGGTGATC |
| *mPadi4*_rev | CTCGAGTCAGGGCACCATGTGCCACCAC |
| hPADI4_D165A_fw | ATCTTCTGCCATGGCCTGCGAGGATGATGAAG |
| hPADI4_D165A_rev | CATCATCCTCGCAGGCCATGGCAGAAGATTCG |
| hPADI4_D168A_fw | CTGCCATGGACTGCGAGGCTGATGAAGTGCTTGAC |
| hPADI4_D168A_rev | GTCAAGCACTTCATCAGCCTCGCAGTCCATGGCAG |
| hPADI4_D167A_fw | CTTCTGCCATGGACTGCGCGGATGATGAAGTGCTTGAC |
| hPADI4_D167A_rev | GTCAAGCACTTCATCATCCGCGCAGTCCATGGCAGAAG |
| hPADI4_N373A_fw | TTCGACTCTCCAAGGGCCAGAGGCCTGAAGG |
| hPADI4_N373A_rev | CTCCTTCAGGCCTCTGGCCCTTGGAGAGTCG |
| hPADI4_R394A_fw | GATTTTGGCTATGTAACTGCAGGGCCCCAAACAGGGGG |
| hPADI4_R394A_rev | CCCCCTGTTTGGGGCCCTGCAGTTACATAGCCAAAATC |
| hPADI4_C166F_fw | CTCGAATCTTCTGCCATGGACTCCGAGGATGATGAAGTGCTT G |
| hPADI4_C166F_rev | CAAGCACTTCATCATCCTCGAAGTCCATGGCAGAAGATTCGA G |
| T7g10M.F48 | TAATACGACTCACTATAGGGTTAACTTTAAGAAGGAGATATAC ATATG |
| long_HA_reverse | TTTCCGCCCCCCGTCCTAAGAACCAGAACCAGAACCTGCATA GTCGGGCACGTCGTATGGGTAGCTGCCGCTGCC |
| short_HA_reverse | TTTCCGCCCCCCGTCCTAAGAACCAGAACCAGAACC |
| PAD4_3_F_NNK1 | CTATAGGGTTAACTTTAAGAAGGAGATATACATATGNNKGATC ATCATTATAGGCATCCGAAGTATTGC |
| PAD4_3_F_NNK2 | CTATAGGGTTAACTTTAAGAAGGAGATATACATATGAGGNNK CATCATTATAGGCATCCGAAGTATTGC |
| PAD4_3_F_NNK3 | CTATAGGGTTAACTTTAAGAAGGAGATATACATATGAGGGAT NNKCATTATAGGCATCCGAAGTATTGC |
| PAD4_3_F_NNK4 | CTATAGGGTTAACTTTAAGAAGGAGATATACATATGAGGGAT CATNNKTATAGGCATCCGAAGTATTGC |
| PAD4_3_F_NNK5 | CTATAGGGTTAACTTTAAGAAGGAGATATACATATGAGGGAT CATCATNNKAGGCATCCGAAGTATTGC |
| PAD4_3_R_NNK6 | TATGGGTAGCTGCCGCTGCCGCAATACTTCGGATGMNNATA ATGATGATCCCTCATAT |
| PAD4_3_R_NNK7 | TATGGGTAGCTGCCGCTGCCGCAATACTTCGGMNNCCTATA ATGATGATCCCTCATAT |

| | |
|---|---|
| PAD4_3_R_NNK8 | TATGGGTAGCTGCCGCTGCCGCAATACTTMNNATGCCTATAATGATGATCCCTCATAT |
| PAD4_3_R_NNK9 | TATGGGTAGCTGCCGCTGCCGCAATAMNNCGGATGCCTATAATGATGATCCCTCATAT |
| PAD4_3_R_NNK10 | TATGGGTAGCTGCCGCTGCCGCAMNNCTTCGGATGCCTATAATGATGATCCCTCATAT |
| PADI4_AttB1_F | GGGGACAAGTTTGTA CAAAAAAGCAGGCTTCACCATGGCCCAGGGGACATTGATCCG |
| PADI4_AttB2_R | GGGGACCACTTTGTACAAGAAAGCTGGGTCTCAGGGCACCATGTTCCACC |
| Pu_linker | [5'Phos]CTCCCGCCCCCCGTCC[SP18][SP18][SP18][SP18][SP18]CC[Puromycin] |

**Supplementary Table 5. Macrocyclic peptides used in this study.** All peptides are cyclic (thioether bond between the Acetylated N-terminus and the cysteine side chain) unless otherwise specified. Abbreviations: Ac – N-terminal acetylation; y – D-Tyrosine; Cit – citrulline; Y(OMe) – O-Methyl-Tyrosine; B – β-Ala; K(Z) – Z attached via an amide bond to the lysine side chain where Z could be: bio (biotin), Cl (cl-alkane). Residues modified from the parent sequences are highlighted in bold. *Mass determined by MALDI.

| Peptide Name | Peptide Sequence | Mass Calculated /Da | Mass Observed /Da |
|---|---|---|---|
| PAD4_1 | yFYRIGFWYPNYQC(S-)G-NH$_2$ | 2016.3 | 2015.3* |
| PAD4_2 | yRDHRSPFDGYC(S-)G-NH$_2$ | 1611.7 | 1611.5 |
| PAD4_3 | yRDHHYRHPKYC(S-)G-NH$_2$ | 1771.0 | 1770.1 |
| PAD4_3i | y**DR**HHYRHPKYC(S-)G-NH$_2$ | 1771.0 | 1769.8 |
| PAD4_4 | yHRLIVVIYVC(S-)G-NH$_2$ | 1473.8 | 1473.6 |
| PAD4_5 | yATPWLIVVLC(S-)G-NH$_2$ | 1373.7 | 1372.8 |
| PAD4_6 | yLVLTIRLVLC(S-)G-NH$_2$ | 1401.8 | 1400.9 |
| PAD4_7 | yYPKGSWGYKLFC(S-)G-NH$_2$ | 1708.0 | 1707.5 |
| PAD4_8 | yTLWTVLVVIC(S-)G-NH$_2$ | 1406.7 | 1427.5* (+Na) |
| PAD4_9 | yAQWYIWVLLC(S-)G-NH$_2$ | 1553.9 | 1553.4 |
| PAD4_10 | yPWEISVWLLYC(S-)G-NH$_2$ | 1668.0 | 1666.3 |
| PAD4_11 | YESC(S-)RYRQVLQL-NH$_2$ | 1596.8 | 1596.0 |
| PAD4_11_i | Y**SE**C(S-)RYRQVLQL-NH$_2$ | 1596.8 | 1597.6 |
| PAD4_12 | YEPC(S-)RFREILDL-NH$_2$ | 1592.8 | 1593.5 |
| PAD4_3_R2A | y**A**DHHYRHPKYC(S-)G-NH$_2$ | 1685.8 | 1685.8 |
| PAD4_3_R7A | yRDHHY**A**HPKYC(S-)G-NH$_2$ | 1685.8 | 1684.1 |
| PAD4_3_R2cit | y**Cit**DHHYRHPKYC(S-)G-NH$_2$ | 1771.9 | 1772.0 |
| PAD4_3_R2K | y**K**DHHYRHPKYC(S-)G-NH$_2$ | 1742.3 | 1742.8 |
| PAD4_3_y1(OMe)Y | Y(OMe)RDHHYRHPKYC(S-)G-NH$_2$ | 1785.0 | 1784.9 |
| P4_3_H4R | yRD**R**HYRHPKYC(S-)G-NH$_2$ | 1790.0 | 1789.4 |
| P4_3_K10I | yRDHHYRHP**I**YC(S-)G-NH$_2$ | 1755.9 | 1755.6 |
| P4_3_K10V | yRDHHYRHP**V**YC(S-)G-NH$_2$ | 1741.9 | 1741.6 |
| P4_3_H4R_K10I | yRD**R**HYRHP**I**YC(S-)G-NH$_2$ | 1775.0 | 1774.9 |
| P4_3_R7N | yRDHHY**N**HPKYC(S-)G-NH$_2$ | 1728.9 | 1728.6 |
| P4_3_R7W | yRDHHY**W**HPKYC(S-)G-NH$_2$ | 1801.0 | 1800.9 |
| P4_3_Y6H | yRDHH**H**RHPKYC(S-)G-NH$_2$ | 1744.9 | 1744.7 |
| PAD4_11_Y1A | **A**ESC(S-)RYRQVLQL-NH$_2$ | 1504.7 | 1503.8 |
| PAD4_11_E2A | Y**A**SC(S-)RYRQVLQL-NH$_2$ | 1538.8 | 1539.4 |
| PAD4_11_S3A | YE**A**C(S-)RYRQVLQL-NH$_2$ | 1580.8 | 1581.3 |
| PAD4_11_C4A | Ac-YES**A**RYRQVLQL-NH$_2$ | 1524.7 | 1565.5 |
| PAD4_11_R5A | YESC(S-)**A**YRQVLQL-NH$_2$ | 1511.7 | 1510.7 |
| PAD4_11_Y6A | YESC(S-)R**A**RQVLQL-NH$_2$ | 1504.7 | 1505.8 |
| PAD4_11_R7A | YESC(S-)RY**A**QVLQL-NH$_2$ | 1511.7 | 1512.2 |
| PAD4_11_Q8A | YESC(S-)RYR**A**VLQL-NH$_2$ | 1539.8 | 1538.8 |
| PAD4_11_V9A | YESC(S-)RYRQ**A**LQL-NH$_2$ | 1568.8 | 1569.4 |
| PAD4_11_L10A | YESC(S-)RYRQV**A**QL-NH$_2$ | 1554.8 | 1555.6 |
| PAD4_11_Q11A | YESC(S-)RYRQVL**A**L-NH$_2$ | 1539.8 | 1540.3 |
| PAD4_11_L12A | YESC(S-)RYRQVLQ**A**-NH$_2$ | 1554.8 | 1555.2 |
| PADI4_11_R5K | YESC(S-)**K**YRQVLQL-NH$_2$ | 1568.3 | 1568.0 |
| PADI4_11_R7K | YESC(S-)RY**K**QVLQL-NH$_2$ | 1568.3 | 1568.4 |

| | | | |
|---|---|---|---|
| PADI4_11_E2Q | YQSC(S-)RYRQVLQL-$NH_2$ | 1593.8 | see 2+ ion 797.8 |
| PADI4_11_E2D | YDSC(S-)RYRQVLQL-$NH_2$ | 1580.8 | 1581.0 |
| PADI4_11B | YESC(S-)Y-$NH_2$ | 858.9 | 858.1 |
| PAD4_3_CAPA | yRDHHYRHPKYC(S-)GSBK(Cl)-$NH_2$ | 2364.4 | 2362.4 |
| PAD4_3i_CAPA | y**DR**HHYRHPKYC(S-)GSBK(Cl)-$NH_2$ | 2364.4 | 2364.3 |
| PAD4_11_CAPA | YESC(S-)RYRQVLQLGSBK(Cl)-$NH_2$ | 2247.5 | 2245.5 |
| PAD4_11i_CAPA | Y**SE**C(S-)RYRQVLQLGSBK(Cl)-$NH_2$ | 2246.9 | 2244.8 |
| P4_7_bio | yYPKGSWGYKLFC(S-)GSBK(Bio)-$NH_2$ | 2220.6 | 2220.1 |
| P4_7scr_bio | yYPKGSWGYKLFC(S-)GSBK(Bio)-$NH_2$ | 2220.6 | 2220.4 |

**Uncropped Western Blots**

Figure S2A

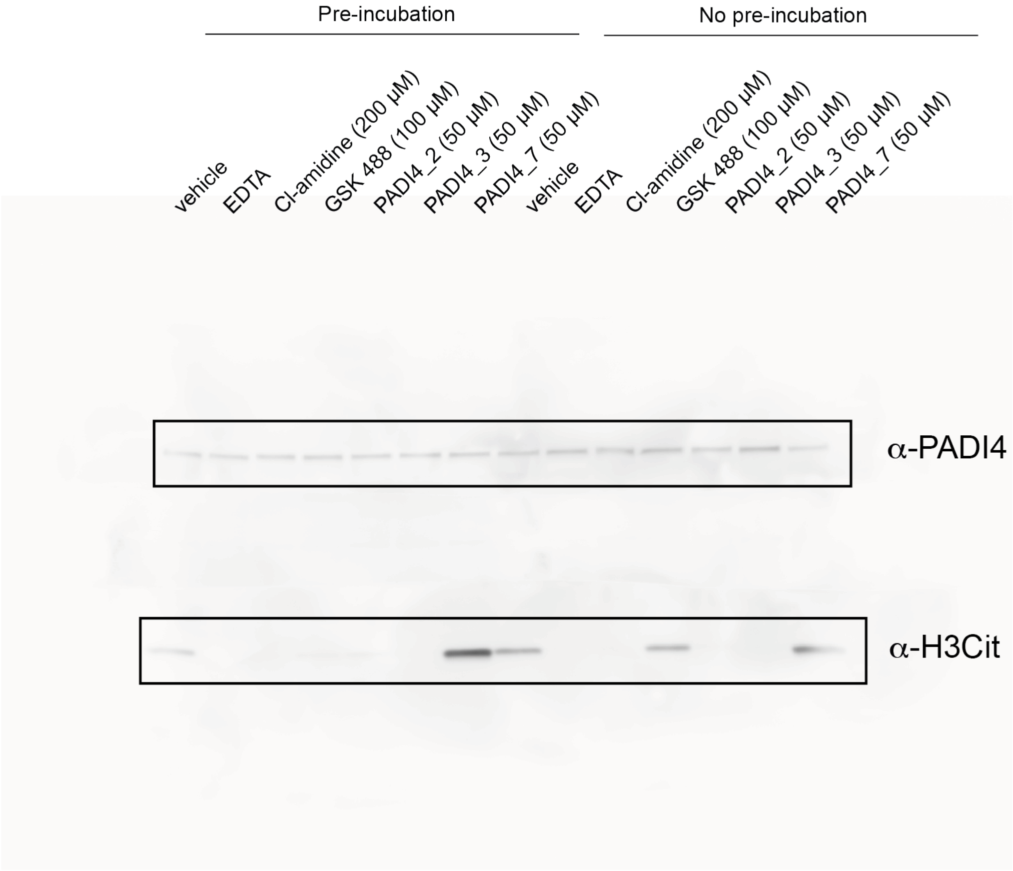

Figure 3D

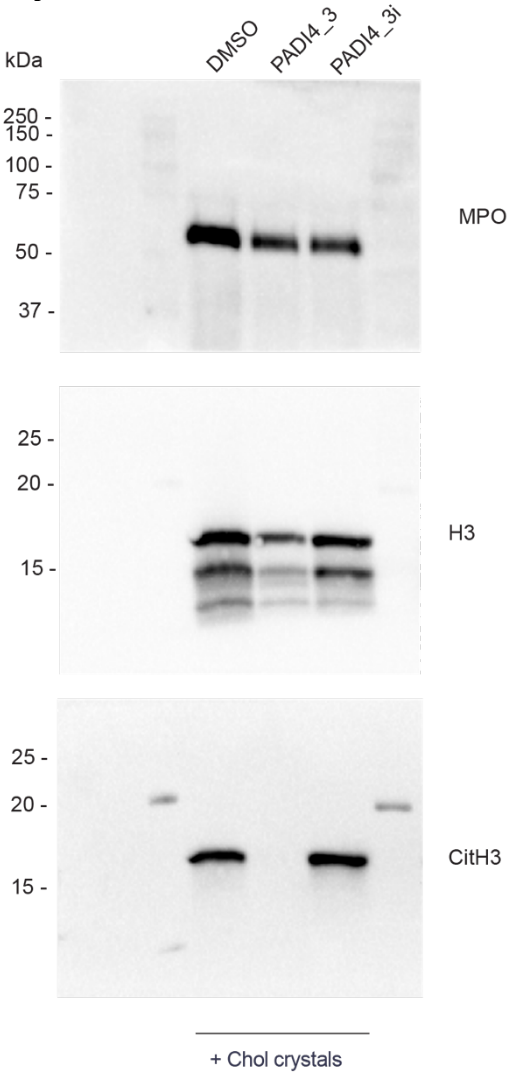
kDa
DMSO
PADI4_3
PADI4_3i
250 -
150 -
100 -
75 -
50 -
37 -
MPO
25 -
20 -
15 -
H3
25 -
20 -
15 -
CitH3
+ Chol crystals

Figure 3F

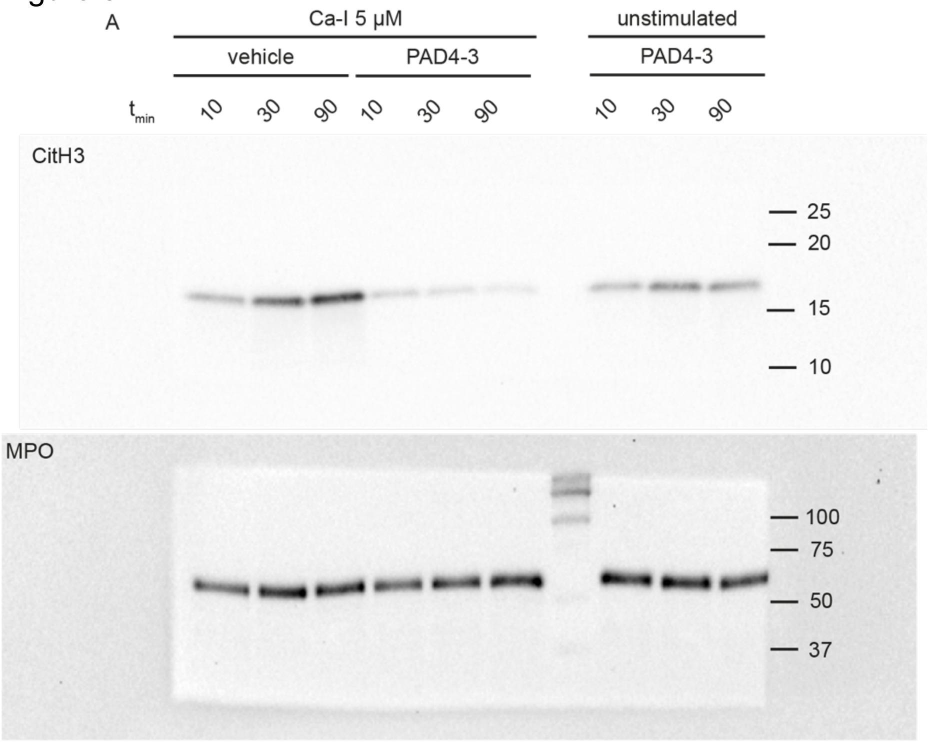

A
Ca-I 5 µM
unstimulated
vehicle
PAD4-3
PAD4-3
t min
10 30 90 10 30 90 10 30 90
CitH3
25
20
15
10
MPO
100
75
50
37
